# Selective pressure by rifampicin modulates mutation rates and evolutionary trajectories of mycobacterial genomes

**DOI:** 10.1101/2023.02.01.525872

**Authors:** E. Cebrián-Sastre, A. Chiner-Oms, R. Torres-Pérez, I. Comas, J.C. Oliveros, J. Blázquez, A. Castañeda-García

## Abstract

Resistance to the frontline antibiotic rifampicin constitutes a challenge to the treatment and control of tuberculosis. Here, we analyzed the mutational landscape of *Mycobacterium smegmatis* during long-term evolution with increasing concentrations of rifampicin, using a mutation accumulation assay combined with whole genome sequencing. Antibiotic treatment enhanced the acquisition of mutations, doubling the genome-wide mutation rate of the wild type cells. While antibiotic exposure led to extinction of almost all wild type lines, the hypermutable phenotype of the Δ*nucS* strain (noncanonical mismatch repair deficient) provided an efficient response to the antibiotic, leading to high rates of survival. This adaptative advantage resulted in the emergence of higher levels of rifampicin resistance, an accelerated acquisition of drug resistance mutations in *rpoB* (β RNA polymerase) and a wider diversity of evolutionary pathways that led to drug resistance. Lastly, this approach revealed a subset of adaptive genes under positive selection with rifampicin that were able to drive novel rifampicin resistance mechanisms.

## Introduction

The acquisition and spread of antimicrobial resistance (AMR) in pathogenic bacteria are major threats to public health. It has been estimated that, without effective healthcare policies to control it, AMR could contribute to 10 million deaths per year by 2050, exceeding the current mortality rate from cancer (*1*). Huge pressure imposed over decades by antimicrobial usage in humans has led to the evolution of genetic variants and the selection and spread of antimicrobial resistant strains.

One of the main ways to achieve antibiotic resistance is through the acquisition of mutations in different chromosomal loci (*2*). Bacterial cells with higher mutation rates (mutators or hypermutators) have a higher probability of acquiring favorable mutations, including those that confer antibiotic resistance. These hypermutable strains, commonly deficient in postreplicative DNA mismatch repair (MMR), can thrive under selective conditions such as antibiotic pressure (*3*).

In addition to driving the selection of resistant variants, many antibiotics can also cause increases in bacterial mutagenesis rates. Bactericidal antimicrobials such as fluoroquinolones, β-lactams, antifolates and aminoglycosides, can induce responses that accelerate mutation rates, driving both the generation of genetic diversity and the selection of resistant variants (*3*–*6*). Production of reactive oxygen species (ROS) and DNA damage responses are central to the accumulation of antibiotic-induced damage, and antibiotic-induced mutagenesis mechanisms can be classified into three largely intertwined groups: induction of the production of ROS, induction of the SOS response and activation of the RpoS-regulated response (*6*).

The mutagenic effect of antibiotics and the selection of hypermutable variants could be especially important for bacterial pathogens such as *Mycobacterium tuberculosis*, which acquire antibiotic resistance exclusively through chromosomal mutations (*7*). Tuberculosis (TB) is one of the top causes of death worldwide and a leading cause of death from a single infectious agent, second only to COVID-19 (*8*). Drug resistance in *M. tuberculosis* makes the control of TB more challenging and remains a major global threat to public health. Close to half a million people develop rifampicin-resistant TB (RR-TB) every year, and 78% of those cases have multidrug resistant TB (MDR-TB), which is resistant to both rifampicin and isoniazid (*9*). The success rate of treatment decreases considerably in cases with drug resistant TB (success rate under 60%, with higher rates of relapse and death), requiring longer treatment with more expensive and toxic second-line drugs (*8*).

Rifampicin is, along with isoniazid, the most powerful first-line antibiotic for TB. It is a bactericidal drug that acts by binding to the DNA-dependent RNA polymerase subunit β (RNAP β), encoded by *rpoB* gene, thereby inhibiting transcription (*10*). Emergence of mutations in the *rpoB* gene represents the main mechanism underlying rifampicin resistance, and is responsible for 90–95 % of rifampicin-resistant *M. tuberculosis* isolates (*11*). The mechanism of resistance in the remaining rifampicin-resistant isolates remains largely unknown, although the involvement of lowered cell permeability and enhanced efflux pumps has been hypothesized (*12*). Interestingly, rifampicin-induced ROS formation has been observed in the pathogen (*13*).

Experimental evolution has been widely used to study antibiotic resistance mechanisms (*14*). The mutation accumulation (MA) assay is a specific type of experimental evolution with single-cell bottlenecks at each serial transfer. MA is designed to allow mutations to occur in a neutral manner without selection, and consequently all non-lethal mutations can be fixed (*15*). When evolving MA cells are exposed to antibiotic selection, drug-induced changes in the rate and molecular landscape of mutations can be quantitatively analyzed without the interference from other factors. Using the MA assay with different bacterial cell lines, combined with whole genome sequencing (WGS), overcomes the limitations of classical methods for determining mutation rates (*16*). Indeed, we have previously used this experimental approach in absence of antibiotic to determine the mutational rate and landscape of a wild type (WT) strain of the surrogate model *Mycobacterium smegmatis* and a noncanonical MMR-deficient Δ*nucS* derivative with a hypermutator phenotype (*17*).

To date, there are few studies that have quantitatively evaluated the bacterial genome-wide mutational profile by MA/WGS after antibiotics exposure (*16*, *18*). In this work, we performed MA/WGS analyses in several independent bacterial lines of *M. smegmatis* wild type and Δ*nucS* strains under increasing concentrations of rifampicin, to characterize the rate and molecular spectrum of genomic mutations and analyze the potential role of hypermutation in response to the antibiotic. In addition, we studied the evolutionary pathways to the acquisition of rifampicin resistance in *M. smegmatis*, with a special emphasis on the study of the appearance of mutations in *rpoB* and the corresponding resistance level conferred over time. We also searched for candidate genes enriched with mutations that may be part of the adaptative response and/or alternative resistance mechanisms. Our results provide quantitative insight into the relationship between antibiotic selective pressure and the rate and molecular spectrum of mutations, the degree to which noncanonical MMR could be involved, and the extent to which elevated mutation rates accelerate the acquisition of drug resistance in mycobacteria.

## Results

### MA experiments under rifampicin selection

In a recent work, we determined mutation rates and mutational spectra of *M. smegmatis* wild type and *nucS*-null strains by a MA assay in the absence of antibiotic (*17*). The experimental design consisted of a parallel evolution for 50 weeks with 11 independent lineages derived from each parental strain, allowing a random accumulation of genomic mutations in the evolved cell lineages. Once mutation rates without antibiotic have been characterized in depth, it is key to analyze the effect of antibiotic selective pressure in the mutational landscape of mycobacteria. It is also interesting to decipher whether noncanonical MMR inactivation could promote the adaptation of mycobacteria to antibiotic stress.

In this study, we have evaluated the effect of the frontline antibiotic rifampicin on mycobacterial evolution by MA experiments and WGS using *M. smegmatis* as a model (wild type and Δ*nucS* strains) (see Materials and Methods). Firstly, the minimal inhibitory concentration (MIC) of rifampicin was determined for both strains in solid media. The parental strains showed identical susceptibility to the drug, each with a MIC value of 2 μg ml^−1^. The MA assay under antibiotic pressure was performed following a similar procedure to that in the absence of antibiotic, but rifampicin concentrations were steadily increased, from 0.25 μg ml^−1^ to 32 μg ml^−1^ throughout the experiment, doubling every 5 weeks (Figure 1). Experimental evolution was carried out with a total of 20 lines derived from each *M. smegmatis* parental strain (wild type and Δ*nucS*) for 40 weeks, by transferring a single random colony to a new plate every 7 days (Figure 1). This strict bottleneck strongly restricted selection, ensuring that each emerged mutation was retained.

**Figure 1.**
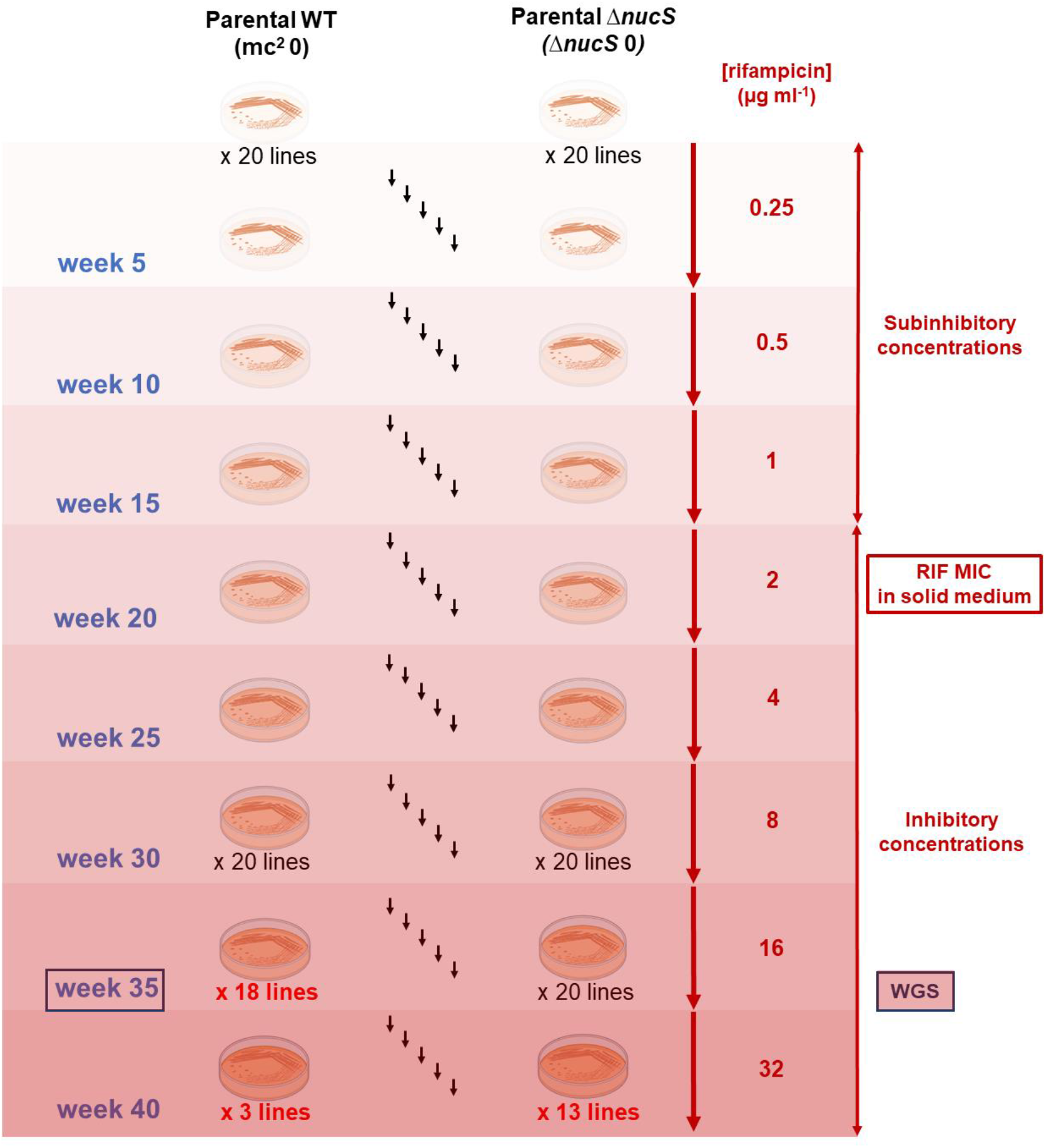
MA experimental evolution with increasing concentrations of rifampicin. The MA assay comprised 40 independent lines, generated from *M. smegmatis* wild type and its *nucS*-null derivative (20 each), and evolved in parallel with increasing antibiotic concentrations. The experimental evolution was carried out for 40 weeks, from 0.25 μg ml^−1^ rifampicin (subinhibitory concentration) until 32 μg ml^−1^ rifampicin (inhibitory concentration), doubling the concentration every 5 weeks. MA lines able to grow at week 35 with 16 μg ml^−1^ rifampicin (18 wild type-derived and 20 Δ*nucS*-derived) were sequenced by WGS. The figure also shows the number of lines that survived at each rifampicin concentration (below the plates) and the number of passages (black arrows). Created with BioRender.com.

At the end of the experiment, by week 40, only 3 of 20 evolved wild type lines were able to survive to the highest rifampicin concentration (32 μg ml^−1^), while the majority of the Δ*nucS* lines (13 of 20), survived. Most of the MA lines (18 of 20 wild type and all 20 Δ*nucS* lines) survived by week 35 in the presence of up to16 μg ml^−1^ rifampicin (Figure 1). The extinction of 85% of the wild type lines but only 35% of *nucS*-deficient lines upon exposure to rifampicin highlights the importance of hypermutability conferred by noncanonical MMR deficiency for resistance to rifampicin in *M. smegmatis*.

### Mutation rates and mutational signatures under antibiotic pressure

Mutation rates in the evolved wild type and Δ*nucS* lines were analyzed by WGS to detect the total number of mutations in each strain. The 18 MA wild type lines and 20 MA Δ*nucS* lines that were able to grow with 16 μg ml^−1^ rifampicin by week 35 were selected for mutational analysis. This antibiotic value is a critical point that allowed us to obtain a full set of WGS data of both strains (beyond that concentration, only three wild type lines survived). Genomic sequences generated by WGS were compared to the *M. smegmatis* mc^2^ 155 reference genome (NC_018289.1) to filter all base pair substitutions (BPSs) and small insertion/deletions (indels) found in each MA line (see Materials and Methods).

Mutation rates per genome (or nucleotide) per generation (Table 1 and Supplementary Tables S1 and S2) were determined by measuring the total number of mutations in the whole genome in relation to the number of cell divisions or generations of each line (see Materials and Methods). WGS analysis identified 148 mutations in MA wild type lines, with 0.009 mutations per genome per division (13.47 x 10^−10^ mutations per nucleotide per generation). A total number of 3181 mutations were detected in Δ*nucS* lines, corresponding to 0.174 mutations per genome per division (258.15 x 10^−10^ mutations per nucleotide per generation). Therefore, the inactivation of noncanonical MMR in the Δ*nucS* lines prompted a ~19-fold increase in the mutation rate of *M. smegmatis* under rifampicin selection.

**Table 1.** Rates of mutation identified in the MA experiment with rifampicin.

| mc<sup>2</sup> 155 (WT) |  |  |  | *ΔnucS* |  |  |
| --- | --- | --- | --- | --- | --- | --- |
|  | No. lines | Total generations | Generations per line | No. lines | Total generations | Generations per line |
|  | 18 | 16,308 | 906 | 20 | 18,315 | 916 |
| Mutation type | No. mutations | Mutation rate per genome per generation (x10<sup>-3</sup>)* | Mutation rate per nucleotide per generation (x10<sup>-10</sup>)* | No. mutations | Mutation rate per genome per generation (x10<sup>-3</sup>)* | Mutation rate per nucleotide per generation (x10<sup>-10</sup>)* |
| **Total** | 148 | 9.07 ± 1.35 | 13.47 ± 2.00 | 3,181 | 173.73 ± 10.69 | 258.15 ± 15.95 |
| **BPSs** | 100 | 6.13 ± 1.35 | 9.11 ± 2.01 | 3,141 | 171.55 ± 10.54 | 254.93 ± 15.73 |
| **Indels** | 48 | 2.94 ± 0.70 | 4.36 ± 1.03 | 40 | 2.17 ± 0.68 | 3.23 ± 1.00 |

\* The mutation rate per genome or nucleotide per generation ± 95% CI (confidence intervals) is shown.

A total of 100 base pair substitutions (BPSs) and 48 small insertions or deletions (indels) were detected in the evolved wild type lines, corresponding to 9.11 x 10^−10^ BPSs and 4.36 x 10^−10^ indels per nucleotide per generation (Table 1 and Supplementary Table S1). The mutational signature reflects that while the majority of mutations were BPSs (with more transitions than transversions), one third of all detected mutations were indels in the wild type lines (Figure 2).

**Figure 2.**
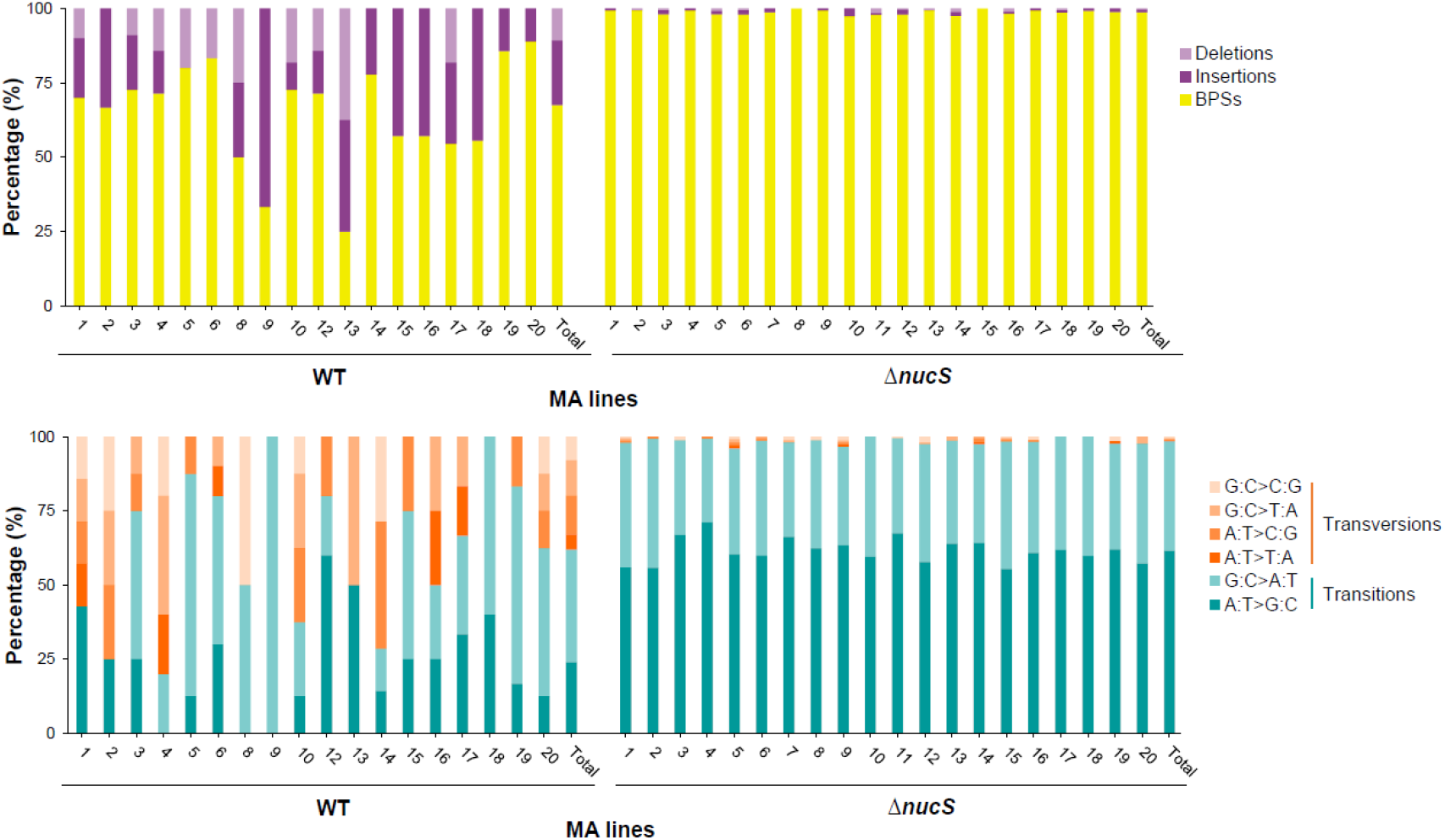
Mutational signature of the MA lines evolved with rifampicin. **A.** Proportion of BPSs, deletions and insertions in the MA lines (wild type and Δ*nucS*). **B.** Relative proportion of the six types of BPSs in the MA lines (wild type and Δ*nucS*). Percentages were calculated with respect to the total number of mutations (A) or BPSs (B) obtained by WGS. Bars are divided in portions with different colors according to the type of mutation, as follows: BPSs (yellow), indels (purple tones), transitions (greeny blue tones) and transversions (orange tones).

In the evolved Δ*nucS* lines, the number of identified BPSs was very high with a total of 3141 BPSs, corresponding to 254.93 x 10^−10^ mutations per nucleotide per generation, while only 40 indels, corresponding to 3.23 x 10^−10^ indels per nucleotide per generation, were detected (Table 1 and Supplementary Table S2). BPSs (almost all transitions) vastly outnumbered indels in evolved Δ*nucS* lines (Figure 2), driving the strong increase in the global mutation rate found in Δ*nucS* genomes, as previously observed during drug-free evolution (*17*). Furthermore, across the Δ*nucS* lines tested, the BPS rate was 28-fold higher compared with the wild type one under rifampicin selection, while no significant change was observed in indel rate. All these results together revealed that the rise in the mutation rate generated by DNA repair deficiency in the Δ*nucS* strain occurred similarly and with the same mutational pattern during drug-free evolution (*17*) and in the evolution under selective antibiotic pressure.

### Rifampicin effect on mutation rates: a comparison in the presence and absence of antibiotic

To determine the effect of rifampicin on the overall mutation rates in this study, we compared the evolution of *M. smegmatis* MA lines with and without (*17*) selective pressure (Figure 3 and 4). A complete dataset of the WGS results (with and without antibiotic) are included in the Supplementary Tables S1 to S5.

**Figure 3.**
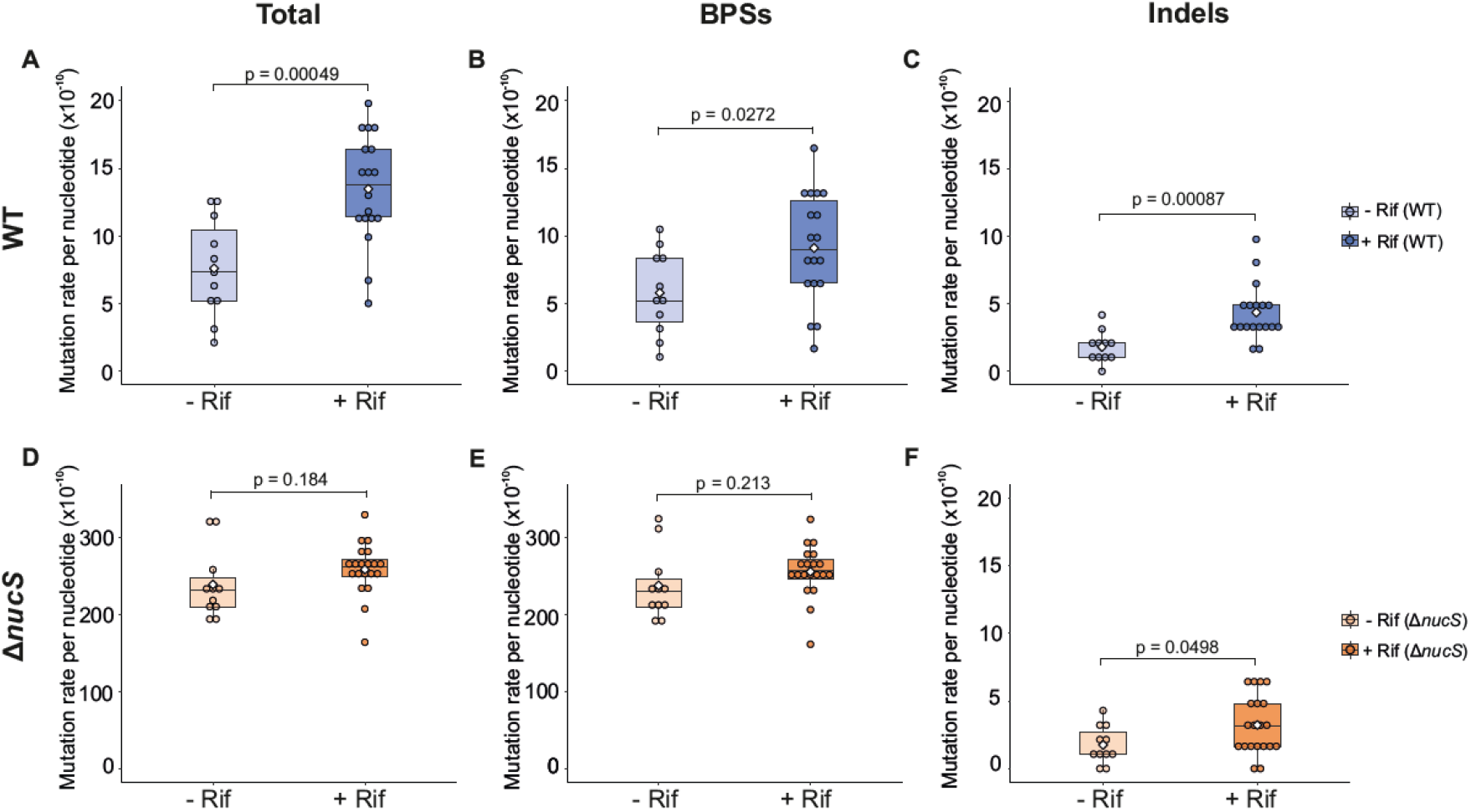
Mutation rates (total mutations, BPSs and indels) in the presence and absence of rifampicin. Boxplots showing the comparison between mutation rates of MA lines evolved with rifampicin (+Rif) (this study) and without antibiotic (-Rif) (*17*). The total mutations, BPSs and indels rates are shown for wild type (**A-C**) (-Rif: light blue, +Rif: dark blue) and Δ*nucS* (**D-F**) (-Rif: light orange, +Rif: dark orange) strains, respectively. Dots represent the mutation rate of each independent MA line. The mean of each mutation rate is indicated with a white diamond. Statistical analyses (t-test) were performed (see Table S5).

**Figure 4.**
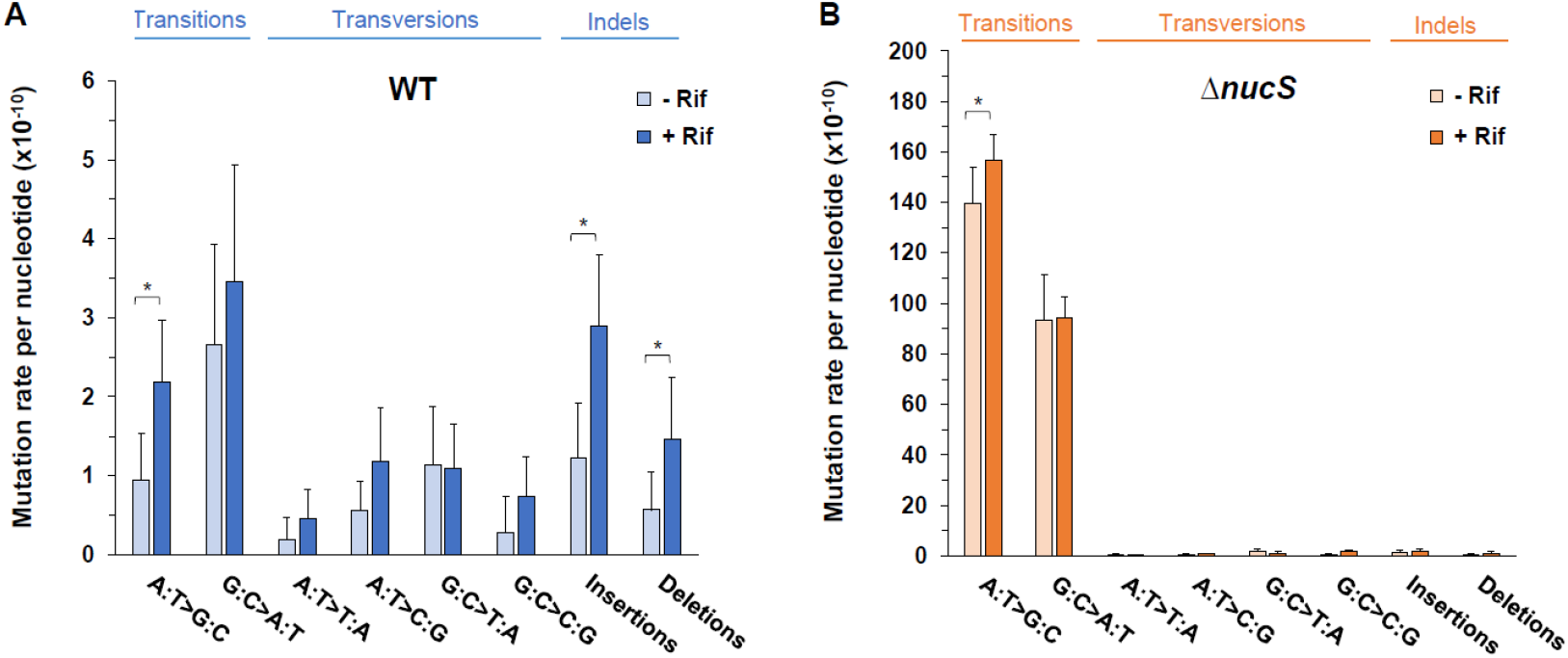
Comparison of the mutational spectra of the MA lines evolved in the presence and absence of rifampicin. **A.** Mutational spectra for *M. smegmatis* wild type lines evolved with rifampicin (dark blue) and without antibiotic (light blue). **B.** Mutational spectra for *M. smegmatis* Δ*nucS* lines evolved in the presence (dark orange) and absence of rifampicin (light orange). Bars represent the mutation rate per nucleotide per generation for each type of DNA mutation obtained by WGS data of MA experiments from this study and our previous work (*17*). Error bars indicate 95% CI (confidence intervals). Statistical analyses (t-test) were performed (see Table S5) (*: p<0.05).

Antibiotic selection led to a doubling in the mutation rate of wild type strain (mutation rate per genome per generation 0.009 with rifampicin versus 0.005 without; mutation rate per nucleotide per generation 13.47 x 10^−10^ versus 7.58 x 10^−10^, respectively) (Figure 3A). No significant effect on the already high mutation rate of Δ*nucS* strain was found in the evolved *nucS*-deficient strain with rifampicin (mutation rate per genome per generation 0.174 with rifampicin versus 0.166 without; mutation rate per nucleotide per generation 258.15 x 10^−10^ versus 238.53 x 10^−10^, respectively) (Figure 3D).

To dissect how rifampicin could promote the emergence of mutations across the genomes, we analyzed the resulting mutational spectrum after experimental evolution in the presence and absence of rifampicin (Figure 3, Figure 4 and Supplementary Tables S1 to S5). In the wild type lines, the rise in the mutation rate was generated by an increase in both BPS and indel rates (Figure 3B and 3C). Although rates of almost all types of mutations were higher in the wild type lines grown with rifampicin, only A:T>G:C transition, insertions, and deletions were significantly increased (2.3-, 2.1-, and 3.7-fold higher with rifampicin than without rifampicin, respectively) (Figure 4A). The sum of all these mutations represents almost half of the total mutations detected in the wild type lines evolved with rifampicin, but less than one third under antibiotic-free evolution. In Δ*nucS* strain, overall mutation rate was not significantly increased under rifampicin selection (Fig 3D), although BPS and indel rates were slightly higher (Figure 3E and 3F). In this sense, the very high mutation rate of this strain could mask any further effect of the antibiotic. Nevertheless, in Δ*nucS* lines, rifampicin selection resulted in one-tenth increase in the A:T>G:C transition, a type of mutation specially increased by *nucS* inactivation (Figure 4B). A moderate increase for insertions and deletions in Δ*nucS* lines was also found (Supplementary Tables S1 to S5), although the contribution of indels to the overall Δ*nucS* mutation rate was very small (Figure 4B).

In summary, this study showed that rifampicin selective pressure had an effect on mutation rates of *M. smegmatis* wild type. Although the DNA repair deficiency due to *nucS* inactivation could have made the Δ*nucS* lines more susceptible to rifampicin-induced mutations, increased acquisition of mutations under rifampicin selection had a major effect on the mutation rates in the wild type lines.

### Evolution of the levels of rifampicin resistance under antibiotic selective pressure

MA evolution allowed us to investigate the evolutionary trajectories of antibiotic resistance following selection by increasing concentrations of rifampicin. The differential response of wild type and Δ*nucS* strains to rifampicin was also elucidated by analyzing the different levels of drug resistance and the appearance of rifampicin resistance mutations in each step of the evolution with antibiotic.

To understand how the drug resistance levels of the MA lines evolved to acquire rifampicin resistance, the susceptibility to rifampicin of all the MA lines obtained in this study was measured. Firstly, we determined MIC values of rifampicin for each MA line at weeks 0, 10, 20, 25, 30, 35 and 40 (see Materials and Methods). Both wild type and Δ*nucS* parental lines were slightly more susceptible to rifampicin in liquid cultures (MIC in 7H9 broth was 1 μg ml^−1^ for both parental strains) than in agar plates (MIC was 2 μg ml^−1^ in 7H10 agar for both strains).

Analysis of the gradual increase in rifampicin resistance during MA evolution is shown in Figure 5. Three levels of drug resistance were defined according to the ranges of rifampicin MIC values: low resistance (MIC of 4–16 μg ml^−1^), intermediate resistance (MIC of 32–128 μg ml^−1^) and high resistance (MIC of 256–1024 μg ml^−1^). When both the wild type and Δ*nucS* lines were exposed to subinhibitory concentrations of rifampicin (< 2 μg ml^−1^), evolved MA lines did not acquire rifampicin resistance and MICs values remained similar to those observed in parental strains (Figure 5). In both strains, the first lines that acquired rifampicin resistance (with a significant increase in MIC) emerged at week 25, under inhibitory concentrations of antibiotic (≥ 2 μg ml^−1^) (Figure 5). From this point forward, under increasing selective pressure, the proportion of resistant lines increased progressively in both strains, even though the proportion of resistant lines (at weeks 25 and 30) was higher in Δ*nucS* lines than in wild type lines (Figure 5). When the resistance levels of the MA lines were examined in both strains, we found a differential response of each strain to the antibiotic pressure (Figure 5). During the final stages of evolution (week 30 onwards), Δ*nucS* lines were able to acquire intermediate to high levels of rifampicin resistance and the majority of highly resistant lines were among the surviving lines by the end of the evolution. By contrast, wild type lines acquired low levels of rifampicin resistance and only a few lines survived and were able to reach intermediate levels of antibiotic resistance by the end of experimental evolution.

**Figure 5.**
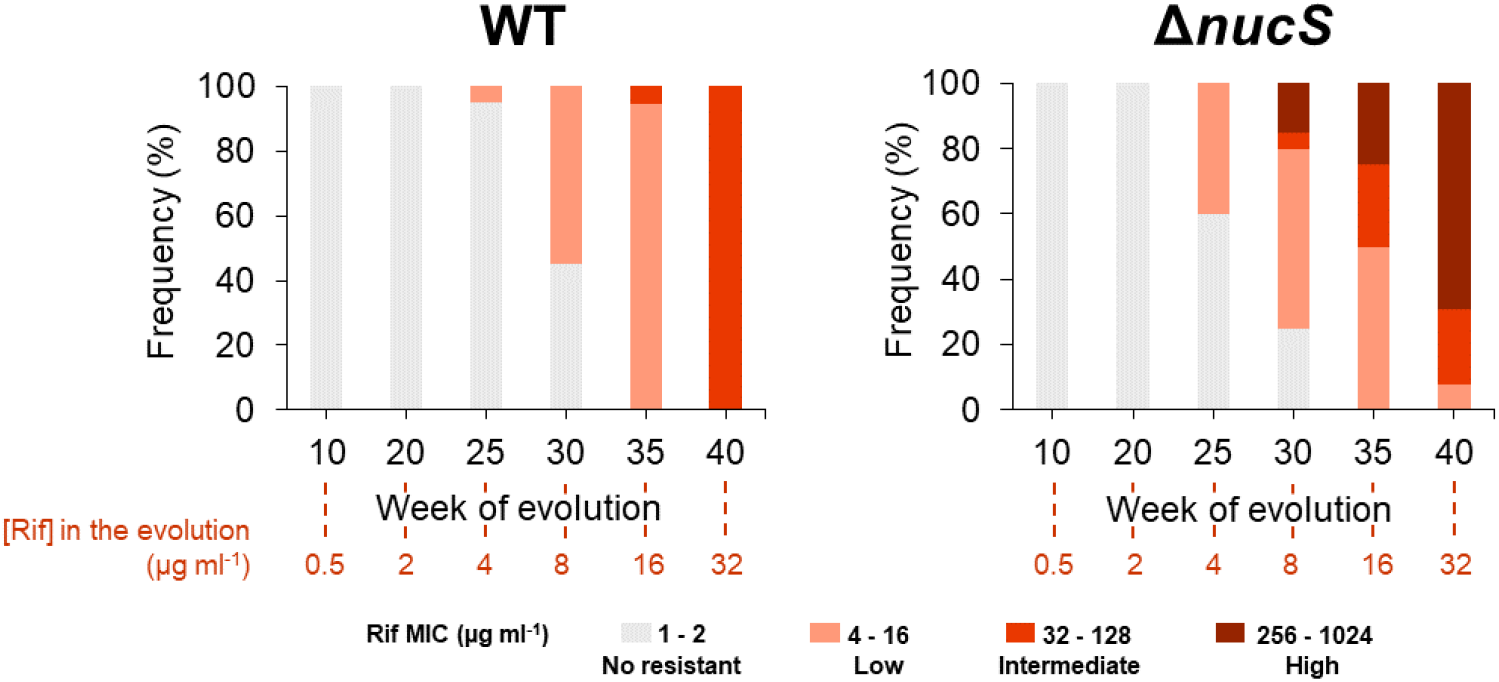
Levels of resistance to rifampicin in the MA lines during experimental evolution under antibiotic selection. The frequency (percentage) of wild type (left) and Δ*nucS* (right) lines with different levels of resistance to rifampicin through the time (in weeks) is shown. The MICs values used to define the levels of rifampicin resistance were the following ones: no resistance, 1 – 2 μg ml^−1^ (grey); low, 4 – 16 μg ml^− 1^ (light red); intermediate, 32 – 128 μg ml^−1^ (medium red); and high, 256 – 1024 μg ml^−1^ (dark red). The numbers shown in red below the graphs indicate the rifampicin concentration used at the corresponding week.

### Acquisition of *rpoB* mutations in wild type and Δ*nucS* lines under rifampicin selection

The highly conserved *rpoB* gene encodes for the beta subunit of the bacterial DNA-dependent RNA polymerase (RNAP β), the main target of rifampicin (*10*). The appearance of mutations in this gene, most of them at the rifampicin-resistant determining region (RRDR), constitutes the main cause of rifampicin resistance in bacteria, including mycobacteria (*19*, *20*). Considering the important role of *rpoB* mutations in the acquisition of rifampicin resistance, we analyzed the *rpoB* sequence from all the MA lines (weeks 0, 10, 20, 25, 30, 35 and 40) during their evolution, in order to assess correlations with the development of antimicrobial resistance. Once *rpoB* mutations were detected, the identity, type and time of appearance of each *rpoB* mutation were characterized (see Materials and Methods).

At the end of experimental evolution, a total of 20 point mutations were detected in *rpoB*, with three in the wild type-derived lines and seventeen in the Δ*nucS*-derived lines (Figure 6A and Supplementary Tables S6 and S7). These data suggest that hypermutability (conferred by noncanonical mismatch repair deficiency) facilitated the acquisition and fixation of a higher number of *rpoB* mutations in Δ*nucS* lines when compared with wild type-derived lines during experimental evolution under antibiotic pressure. We observed one transition and two transversions in *rpoB* in the wild type-derived lines, while all mutations found in *rpoB* of Δ*nucS* lines were transitions, demonstrating that inactivation of *nucS* led to an increase in the number of transitions (Supplementary Tables S6 and S7) (*17*, *21*).

**Figure 6.**
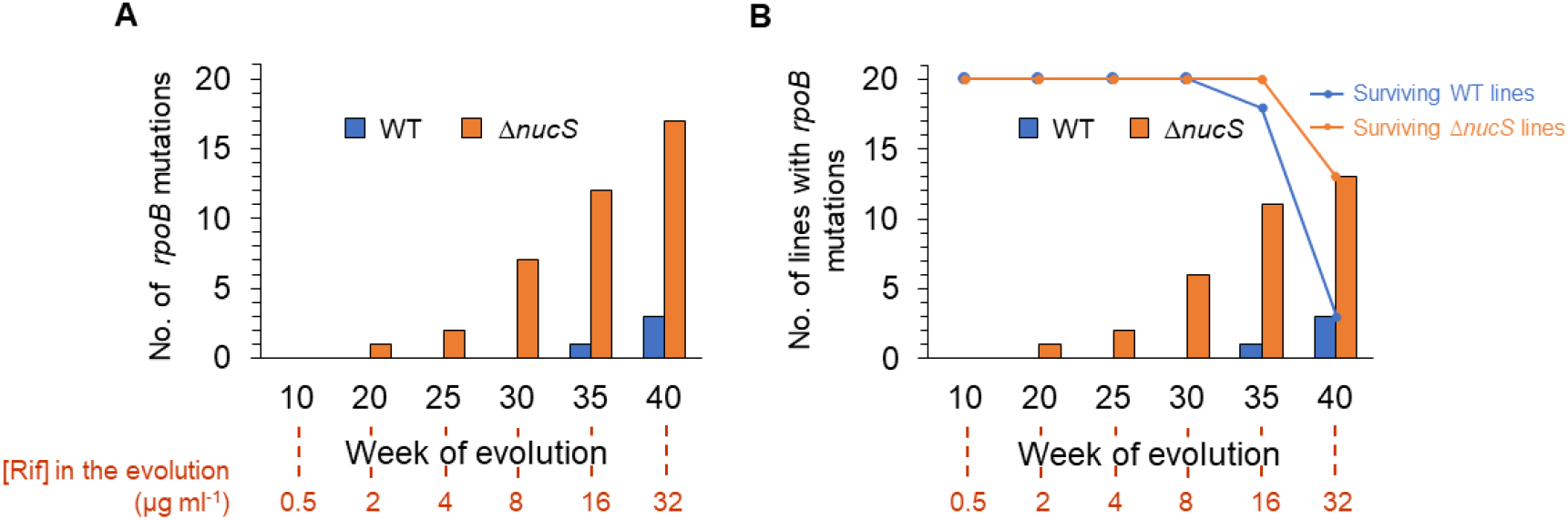
Acquisition of *rpoB* mutations in the MA lines evolved with rifampicin. **A.** Dynamics of the total number of *rpoB* mutations in the MA experiment. Bars show the total number of *rpoB* mutations that accumulated all the MA lines through the time. **B.** Dynamics of the total number of lines with *rpoB* mutations in the MA experiment. Bars represent the number of lines containing *rpoB* mutations though the time (in weeks). Dots of the line plot represent the number of surviving lines (wild type, blue; Δ*nucS*, orange). The numbers shown in red below the graphs indicate the rifampicin concentration used at the corresponding week.

When the dynamics of *rpoB* mutation appearance was assessed (Figure 6A), we found that all *rpoB* mutations arose under inhibitory concentrations of rifampicin and most of them during the final steps of the evolution when the cell lines were exposed to high concentrations of antibiotic. While the wild type strain did not acquire any *rpoB* mutations until week 35 (16 μg ml^−1^ rifampicin), *rpoB* mutations appeared earlier in the hypermutable *nucS*-deficient strain, with the first detected at weeks 20-25, one third arising by week 30, and then increasing through weeks 35-40. Therefore, the absence of *nucS* favored a more rapid acquisition of *rpoB* mutations during experimental evolution under rifampicin selection.

A strong association between the acquisition of *rpoB* mutations and enhanced survival on rifampicin was observed (Figure 6B). Only evolved lines harboring *rpoB* mutations survived under selection with 32 μg ml^−1^ rifampicin at the end of the evolution (week 40). As a result, *rpoB* mutations were observed in all the three wild type-derived lines, while all the remaining thirteen Δ*nucS*-derived lines also had mutations in *rpoB*. Only two Δ*nucS* lines harboring *rpoB* mutation did not survive until week 40.

### Analysis of the levels of rifampicin resistance conferred by *rpoB* mutations

Analysis of the identity of the emerged *rpoB* mutations detected a total of sixteen different mutated positions (codons), fifteen nonsynonymous mutations and one synonymous mutation. As a result, a wide diversity of amino acids substitutions was generated in the RNAP β protein sequence in the evolved lines, with three and twelve different types of amino acid substitutions observed in the wild type and Δ*nucS* strains, respectively (Figure 7A).

**Figure 7.**
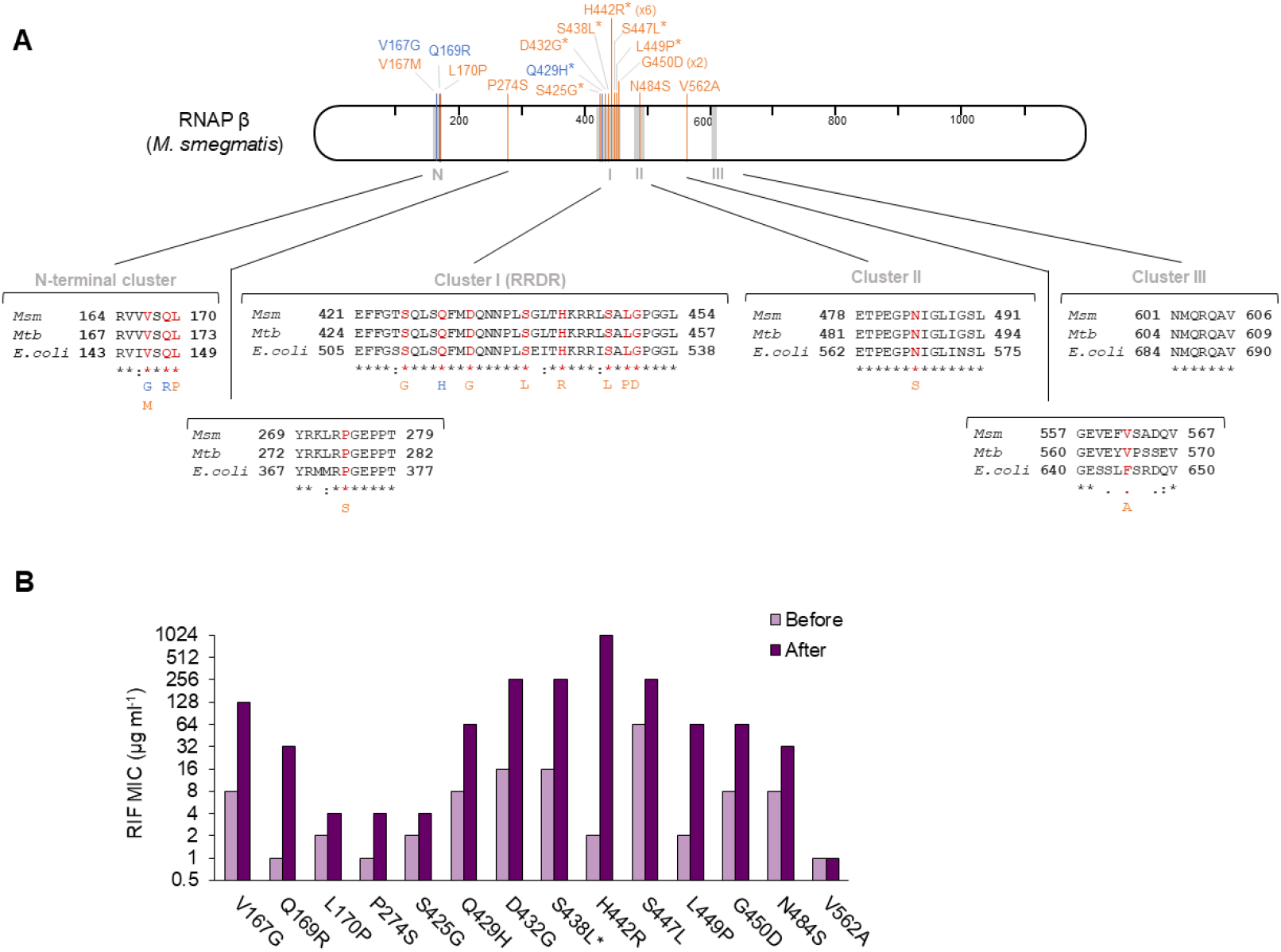
Mutations in *rpoB* in the MA lines: identification and effect on antibiotic resistance. **A.** Types of RNAP β substitutions generated by *rpoB* mutations. *M. smegmatis* RNAP β is represented with amino acid numbering. Gray regions indicate the rifampicin resistance clusters described in *E. coli* (*20*, *23*). Amino acid substitutions generated by *rpoB* mutations in the MA lines are shown in blue (wild type lines) and orange (Δ*nucS* lines). Mutations associated with resistance in *M. tuberculosis* by the WHO catalogue (*26*) are marked with asterisks. Below, protein alignment of RNAP β of *M. smegmatis* (*Msm*), *M. tuberculosis* (*Mtb*) and *E. coli*, with the amino acid positions where mutations were detected (red). **B.** Effect of RNAP β substitutions on rifampicin resistance. Bars show the rifampicin MIC of the MA lines before (light purple) and after (dark purple) the acquisition of each *rpoB* mutation. When the *rpoB* mutation was present in more than one line (H442R and G450D), a representative example is shown. *S438L substitution appeared together with V167M at the same week of the evolution.

Four different conserved clusters of RNAP β amino acid substitutions (N-terminal cluster, cluster I/RRDR, cluster II and cluster III) have been identified in rifampicin-resistant isolates (Figure 7A) (*22*, *23*). In this study, most of the RNAP β substitutions found in the evolved lines were located in the RRDR region (cluster I), the main region with rifampicin resistance-conferring mutations in prokaryotes (*20*, *24*). The remaining amino acid substitutions were found scattered over different regions of the RNAP β sequence, including the N-terminal cluster, with four substitutions, and cluster II, with one substitution (Figure 7A). The hypermutator phenotype of the Δ*nucS* strain facilitated the acquisition of a wider range of *rpoB* mutations compared with the wild type one, with several RNAP β substitutions able to diversify the pathways to antibiotic resistance (see next section).

The most common RNAP β amino acid mutation found in the higher number of MA lines (six Δ*nucS* lines) was H442R. This rifampicin resistance mutation corresponds to one of the most frequent and prominent mutations found in *M. tuberculosis* clinical strains (H445R mutation in *M. tuberculosis* RNAP β), conferring a high level of drug resistance (*25*). Some Δ*nucS* lines also contained other well-known rifampicin resistance mutations, D432G, S438L and S447L, all of them among prevalent drug resistance mutations in *M. tuberculosis* clinical strains (D435G, S441L and S450L mutations in *M. tuberculosis* RNAP β) (*25*, *26*). Interestingly, we also observed a “borderline” resistance mutation in a Δ*nucS*-derived line, L449P (L452P mutation in *M. tuberculosis*), conferring intermediate level of rifampicin resistance (*26*, *27*). While Δ*nucS* lines acquired several different high-level rifampicin resistance mutations under antibiotic pressure due to its hypermutator phenotype, only one well-known rifampicin resistance mutation was detected in the RRDR region of a wild type line, namely Q429H (corresponding to Q432H mutation in *M. tuberculosis* RNAP β, a high confidence *M. tuberculosis* resistance mutation according to WHO criteria) (*26*, *28*).

To evaluate the effect of the different *rpoB* mutations on the acquisition of rifampicin resistance, we compared the MIC values of the evolved lines before and after the emergence of each *rpoB* mutation (Figure 7B). Once all *rpoB* mutations were investigated, most *rpoB* mutant lines exhibited increased MIC values for rifampicin associated with the emergence of the corresponding *rpoB* mutation. These results support a strong link between *rpoB* mutations and the acquisition of drug resistance (although the effect of some other simultaneous mutations cannot be ruled out). Most RRDR mutations were found in isolates that showed high level of rifampicin resistance (≥256 μg ml^−1^), while mutations located in clusters outside the RRDR seemed to generate low to intermediate levels of rifampicin resistance (Figure 7B).

### Evolutionary pathways to the acquisition of rifampicin resistance

MA evolution under antibiotic pressure with independent cell lines constitutes a paradigm to investigate the different evolutionary pathways of antibiotic resistance. To reconstruct the evolutionary trajectories to rifampicin resistance, we first focused on the acquisition of rifampicin resistance by *rpoB* mutations and the corresponding MIC values (rifampicin resistance levels) from each isolate during experimental evolution (Figure 8 and Supplementary Tables 6 and 7).

**Figure 8.**
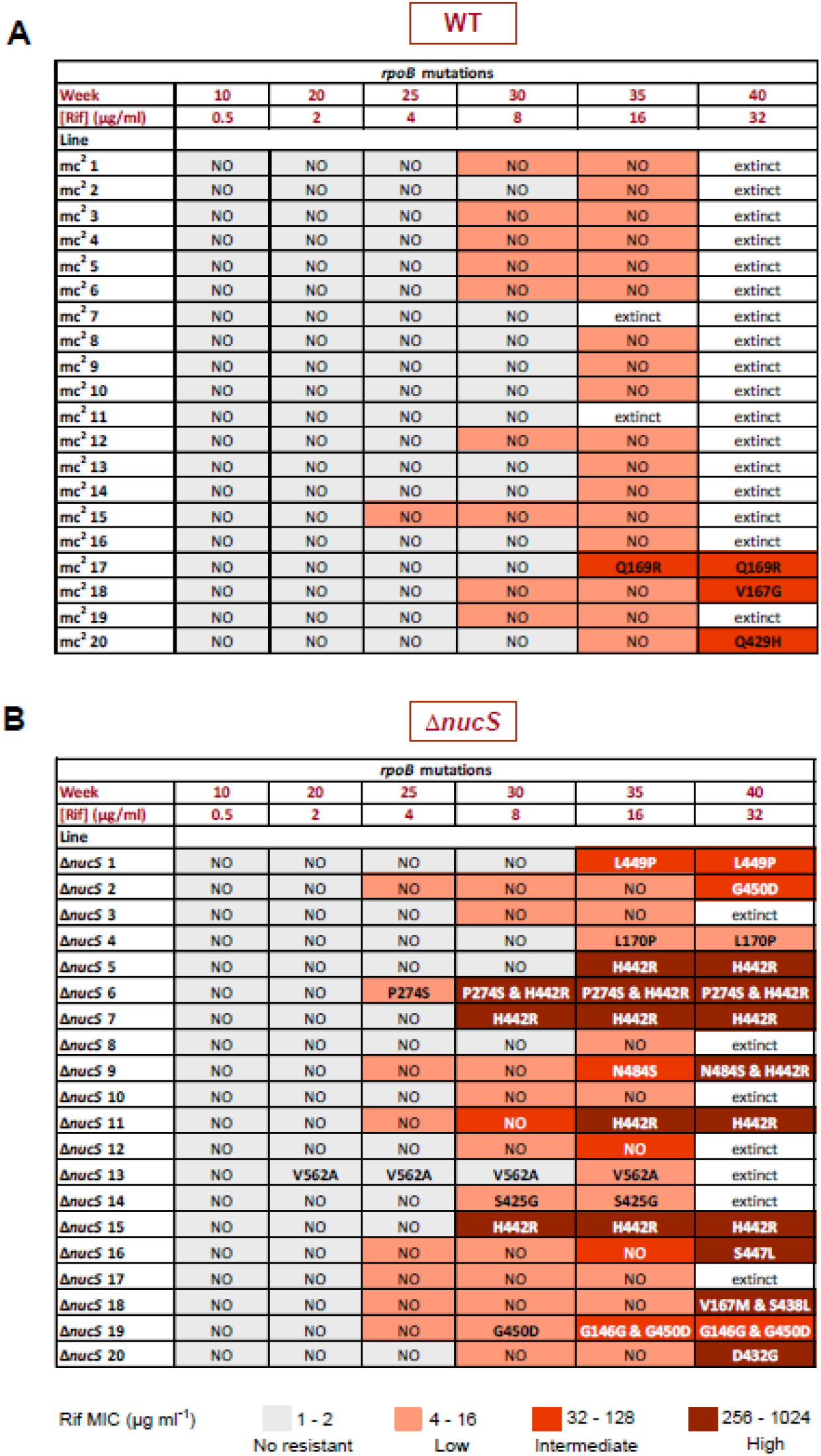
Evolutionary trajectories of the MA lines. **A**. Pathways to acquire rifampicin resistance in the MA wild type lines. **B**. Pathways to acquire rifampicin resistance in the MA Δ*nucS* lines. Tables show the emergence of *rpoB* mutations in each MA line during the experimental evolution (in weeks). Levels of rifampicin resistance of the evolved lines are indicated according to their MICs values, with the following color code: no resistance, 1 – 2 μg ml^−1^ (grey); low, 4 – 16 μg ml^−1^ (light red); intermediate, 32 – 128 μg ml^−1^ (medium red); and high, 256 – 1024 μg ml^−1^ (dark red).

Investigations into the evolutionary pathways leading to rifampicin resistance in wild type-derived lines (Figure 8A) revealed that almost all were able to acquire low levels of drug resistance by *rpoB*-independent mechanisms, with almost half of the wild type lines surviving at week 30 and all the surviving lines at week 35 having low levels of resistance. From week 35, the emergence of *rpoB* mutations led to the acquisition of intermediate levels of drug resistance in three wild type lines, allowing them to grow until the end of the evolution by week 40.

In Δ*nucS*-derived lines (Figure 8B), we observed that the pathways leading to rifampicin resistance were more complex, multiple and diverse than the wild type ones. Δ*nucS* lines reached low levels of resistance earlier, by week 25. Then, most of them acquired intermediate to high rifampicin resistance levels through mutations of *rpoB*, from weeks 25 to 40, leading to a high proportion of surviving drug-resistant isolates.

The most frequent pathway to acquire rifampicin resistance among the Δ*nucS* lines was through RRDR mutations (Figure 8B). The emergence of a RRDR mutation in *rpoB* often conferred high level of drug resistance from no or lower resistance levels (H442R mutation observed in Δ*nucS* lines 5, 7, 11 and 15; D432G mutation in Δ*nucS* line 20; S447L mutation in Δ*nucS* line 16). In some cases, *rpoB* mutations generated intermediate levels of drug resistance in one step, as observed in the L449P mutation in Δ*nucS* line 1 or the G450D mutation in Δ*nucS* line 2. All the Δ*nucS* lines with intermediate or high levels of rifampicin resistance through RRDR mutations were able to survive by the end of the experimental evolution. Alternatively, few Δ*nucS* lines acquired non-RRDR *rpoB* mutation alone (L170P mutation in Δ*nucS* line 4, V562A in Δ*nucS* line 13 and S425G in Δ*nucS* line 14), although these Δ*nucS* lines had low level of resistance and, in some cases, they did not progress further.

Although a single-step progression was the most frequent pathway leading to drug resistance, Δ*nucS* strain exhibited multiple different evolutionary trajectories, including in some cases a multi-step acquisition of rifampicin resistance-conferring mutations (Figure 8B). In two Δ*nucS* lines, Δ*nucS* 6 and 9, we initially detected low to intermediate rifampicin resistance levels by acquisition of a *rpoB* mutation outside the RRDR region and then high levels of drug resistance conferred by the sequential mutation H442R in the RRDR region. Another trajectory found in the Δ*nucS* lines was the simultaneous acquisition of two different *rpoB* mutations, leading to a high level of resistance in the double mutant (Δ*nucS* line 18 with mutations V167M and S438L).

Acquisition of low and even intermediate levels of rifampicin resistance was observed in some cases in the absence of any *rpoB* mutation (Figure 8). A deeper study of the evolution of rifampicin MIC values through the time (see Supplementary Tables 6 and 7) revealed multiple cases where the level of antibiotic resistance increased (4 to 16-fold increase in the MIC values) without any associated *rpoB* mutation. Therefore, mutation of genes other than *rpoB* could contribute to the acquisition of rifampicin resistance in mycobacteria.

### Identification of candidate genes under positive antibiotic selection by rifampicin

Increasing concentrations of antibiotic during experimental evolution should select and fix drug resistance-conferring mutations in specific genes associated with antibiotic resistance. Genetic variants in rifampicin resistance-associated genes should then be subjected to positive selection, leading to a high number of mutations in these genes in the evolved lines. Although rifampicin resistance is mainly associated with *rpoB* mutations, alternative pathways could have a role in a proportion of drug resistant strains.

To provide insights into the complex mechanisms underlying rifampicin resistance, genes that were enriched with mutations during experiment evolution with rifampicin were identified and then compared with the results generated by neutral evolution without antibiotic. We expected to find a higher number of mutations in those genes under antibiotic selection (evolution with rifampicin) in comparison with the accumulation of random mutations (evolution without antibiotic). The use of the Δ*nucS* strain in this experimental approach was a valuable tool to detect loci under selection, as a large number of mutations accumulated across the whole *M. smegmatis* genome during the course of experimental evolution.

All mutations (SNPs and indels) detected in the evolved lines by WGS of both experimental evolution studies, with and without antibiotic, were included in the analysis to filter the genes under selection. Genes that accumulated four or more mutations under antibiotic selection and more than double the number of mutations observed in the absence of antibiotic were considered as candidate genes. According to these criteria, 24 genes were selected, with increased number of allelic variants over the expected random ones (Table 2).

**Table 2.** List of candidate genes under positive selective pressure in the MA experiment with rifampicin.

| Gene ID (NC_018289.1) | Description | No. of mutations |  |
| --- | --- | --- | --- |
|  |  | + Rif <sup>a</sup> | - Rif <sup>b</sup> |
| MSMEI_0106 (*pntA*) | Proton-translocating NAD(P)(+) transhydrogenase | 9 | 1 |
| MSMEI_1105 | Amino acid permease-associated region | 4 | 0 |
| MSMEI_1250 | NERD domain protein | 4 | 1 |
| MSMEI_1328 (*rpoB*) | DNA-directed RNA polymerase subunit beta | 14<sup>c</sup> | 4 |
| MSMEI_1389 (*lldD1*) | L-lactate dehydrogenase (Cytochrome) | 4 | 0 |
| MSMEI_1418 | Putative integral membrane protein | 4 | 0 |
| MSMEI_1810 (*selB*) | Selenocysteine-specific translation elongation factor | 5 | 0 |
| MSMEI_1836 (*mtrB*) | Sensor histidine kinase | 5 | 1 |
| MSMEI_1903 (*mchK*) | Transmembrane cation transporter | 8 | 0 |
| MSMEI_2699 | Nucleic acid binding OB-fold tRNA/helicase-type | 11 | 1 |
| MSMEI_2701 (*trkB*) | Trk system potassium uptake protein | 4 | 0 |
| MSMEI_2763 (*narB*) | Putative oxidoreductase (Nitrate reductase) | 5 | 0 |
| MSMEI_2973 (*carB*) | Carbamoyl-phosphate synthase large chain | 4 | 1 |
| MSMEI_3041 | Probable alpha-(1→6)-mannopyranosyltransferase | 4 | 0 |
| MSMEI_3863 | HAD-superfamily hydrolase subfamily IIB | 4 | 1 |
| MSMEI_4262 | Carbohydrate-binding protein | 4 | 0 |
| MSMEI_4371 (*dnaG*) | DNA primase | 4 | 0 |
| MSMEI_4439 (*ssuD*) | Nitrilotriacetate monooxygenase component A | 4 | 1 |
| MSMEI_4625 (*mmpL*) | Transmembrane transport protein | 4 | 0 |
| MSMEI_5007 (*narH*) | Respiratory nitrate reductase (Beta chain) | 4 | 0 |
| MSMEI_5012 | Glycoside hydrolase, family 3-like protein | 4 | 1 |
| MSMEI_5666 (*purL*) | Phosphoribosylformylglycinamide synthase subunit | 4 | 1 |
| MSMEI_5993 | Carboxymuconolactone decarboxylase | 4 | 1 |
| MSMEI_6055 | Conserved transmembrane protein | 5 | 1 |

<sup>a</sup> Data from the MA experiment with rifampicin (this work).
<sup>b</sup> Data from the MA assay in absence of antibiotic (17).
<sup>c</sup> 13 *rpoB* mutations were detected by WGS and confirmed by PCR, 1 *rpoB* mutation was only identified by PCR.

The loci under selective pressure include multiple genes related to membrane transport, metabolic enzymes (including several oxidoreductases) and some hypothetical conserved proteins with unknown functions (Table 2). *rpoB* is the gene with the highest number of mutations among the selected genes, being under a strong positive selection in the assay (dN/dS = 4.19) (*29*). In summary, the evolution with antibiotic exerted selective pressure on the *M. smegmatis* genome that resulted in the appearance of more mutations in specific genes, with a potential role in adaptation and resistance to rifampicin.

### Acquisition of rifampicin resistance by *rpoB*-independent mechanisms

Although the emergence of high levels of drug resistance depends on *rpoB* mutations, the evolution with rifampicin showed that some evolved lines reached low to intermediate resistance levels in the absence of any *rpoB* mutation (see previous sections). In this case, some genes under positive selection with a higher number of mutations could be genes with drug resistance-conferring mutations, that could explain, in some cases, the acquisition of rifampicin resistance during experimental evolution.

Among the candidates with positive selection (Table 2), we found that mutations in *mchK* and *trkB* genes could be associated with antibiotic resistance during experimental evolution. In this sense, *mchK* and *trkA* (a *trkB* homolog) genes were previously selected in a screening of a library searching for rifampicin resistance genes and led to 8 to 16-fold increase in rifampicin resistance by gene deletion (*30*, *31*). *mchK* and *trkB* genes encode proteins that mediate K^+^ uptake into the cells: MchK is an ion channel protein and TrkB is a regulator of K^+^ uptake (as TrkA, both encoded in the same operon) (*30*, *31*).

To provide insight into the role of ion transport in rifampicin resistance, we detected the emergence of mutations in *mchK* and *trkB* by sequencing (see Materials and Methods), and then analyzed their levels of resistance (Supplementary Figure S1). During experimental evolution, eight MA lines (three wild type lines and five Δ*nucS* lines) with *mchK* mutations and four Δ*nucS* lines with *trkB* mutations were detected. The emergence of most *mchK* and *trkB* mutations were associated with moderate increases in MIC values (2-fold to 8-fold increases) (Supplementary Figure S1). In some cases, they seemed to contribute to the acquisition of low levels of resistance (*mchK* mutations in wild type lines 4, 6 and 19, and *ΔnucS* lines 2, 9 and 19; *trkB* mutation in *ΔnucS* line 20) or even intermediate level of resistance (*trkB* mutation in *ΔnucS* lines 12). The emergence of *mchK* and/or *trkB* mutations was frequently followed by a further mutation in *rpoB* in the *ΔnucS* lines (seven lines in total), leading to a high level of rifampicin resistance in these lines.

These results highlight the effect of alternative resistance mechanisms on the evolution of drug resistance in mycobacteria. A subset of genes under positive pressure could participate in the general response of mycobacteria to antibiotic-induced stress, including in some cases different pathways of drug resistance. In conclusion, our experimental approach could be a useful tool to detect loci and gene variants involved in the evolutionary pathways to antibiotic adaptation and resistance.

## Discussion

MA/WGS experiments have been widely used to measure mutation rates in prokaryotes and eukaryotes (*32*, *33*), as well as to evaluate the effect of antibiotic selection on genome-wide mutations in bacteria (*14*, *16*, *18*). In this work, the effect of long-term exposure to rifampicin was analyzed at the genome-wide level by MA/WGS for the first time in the mycobacterium *M. smegmatis.* Rifampicin treatment resulted in a genome-wide mutagenic effect, with mutation rates duplicated in the wild type strain and a weaker effect on the already hypermutator Δ*nucS* strain. While a two-fold increase in mutation rate could be considered mild, it has been demonstrated that even modest changes in mutation frequencies can significantly influence the evolution of antibiotic resistance (*34*). Therefore, rifampicin exposure acts as a selective agent for potential drug resistance mutations, as well as promoting genetic diversity, thus increasing the likelihood of acquisition and fixation of mutations that confer drug resistance.

Many different antibiotics have been described as promoters of genetic variation through a transient increase in mutation rates and genetic instability (*35*), including some that directly affect DNA replication and genome integrity (*36*, *37*). The inhibitory effect of rifampicin on DNA expression may also contribute to generate genetic instability, as there is a strong connection between DNA replication and transcription, with specific DNA repair mechanisms involved in the resolution of replication /transcription conflicts that affect DNA integrity (*38*, *39*). Treatments with some antibiotics (i.e. β-lactams, aminoglycosides and quinolones) can result in an increased generation of toxic ROS, resulting in higher mutation rates (*5*, *40*) and bacterial death (*41*–*43*). Along with isoniazid and pyrazinamide (*44*), rifampicin induces an oxidative burst that contributes to the antibiotic-mediated killing in *M. tuberculosis* (*13*). Rifampicin may also mediate DNA damage through ROS production in *M. smegmatis*, as this antibiotic affects the ROS levels in *M. smegmatis*, improving the susceptibility to this antibiotic (*45*, *46*) and has a synergic effect with oxidative agents in different mycobacterial models (*47*). Interestingly, enhanced ROS generation depends on the binding of the antibiotic to the RNAP and rifampicin-dependent increases in ROS levels are not observed in resistant strains with *rpoB* mutations (*13*). Evolved Δ*nucS* lines, most of them with *rpoB* mutations, could be desensitized to the mutagenic effect of rifampicin during experimental evolution. Further efforts are needed to identify the mechanisms underlying antibiotic-induced mutagenesis of rifampicin in mycobacteria.

This study investigated the effect of NucS-dependent noncanonical MMR deficiency on the evolution of drug resistance mutations *in vivo* in mycobacteria, and revealed that the Δ*nucS* strain (hypermutator) had a higher probability of acquiring adaptive mutations, including those that confer rifampicin resistance. Hypermutability confers evolutionary advantages for bacterial adaptation (*48*), by favoring the acquisition of mutations that promote the response to stressful and challenging environments such as antibiotic treatments (*49*–*51*). In Δ*nucS* lines, the mutation-driven emergence of antibiotic resistant isolates increased rapidly and strongly, with larger number of drug resistance-conferring mutations (*rpoB* mutations) and higher levels of drug resistance compared with wild type ones. Therefore, inactivation of noncanonical MMR accelerated the appearance and fixation of drug resistance to rifampicin in mycobacteria. Not surprisingly, hypermutators are widespread in clinical isolates of multiple pathogens (*52*–*54*), where the beneficial acquisition of antibiotic resistance seems to be a key factor for their success during infection (*3*, *55*, *56*). The MA experimental approach revealed the mechanisms that could support the potential rise and spread of a *nucS*-deficient strain in a population under antibiotic pressure. In a bacterial population, indirect selection or second order selection (a linkage between a beneficial mutation and the mutator allele), supports the selection for hypermutator clones (*57*, *58*). In this study, the success of the hypermutator Δ*nucS*, with high survival rates under increasing concentration of rifampicin, correlates with the co-selection of high-level resistance mutations in *rpoB*, as all surviving MA lines had at least one *rpoB* mutation.

The high evolvability (ability of an organism to generate genetic variation during evolution) of the *nucS*-deficient strain is a powerful tool for the acquisition of rifampicin resistance during evolution under antibiotic selection. Δ*nucS* lines generated a wider diversity of antibiotic resistance-conferring mutations across the *rpoB* gene than the wild type lines, with single and even double mutations (simultaneous or successive), leading to an increased number and range of evolutionary pathways leading to rifampicin resistance. Since the acquisition of *rpoB* mutations is frequently associated with fitness cost (*59*, *60*) and compensatory mutations (*61*), this study opens the potential to search for new compensatory mutations that may relieve the loss of fitness associated with rifampicin resistance. Although several studies support the emergence and prolonged survival of hypermutators (*62*–*64*), it would be interesting to evaluate the effect of a constitutive (heritable) *nucS*-deficient strain in terms of growth and fitness during long-term experimental evolution.

The analysis of the MA evolution under antibiotic pressure provides details on the mechanisms of acquisition of rifampicin resistance. Most rifampicin resistant *M. tuberculosis* strains have nonsynonymous mutations in the *rpoB* gene (RNAP beta subunit), the majority in the RRDR region (*19*), with amino acid substitutions that prevent the effective binding of rifampicin within the beta subunit (*65*). It is remarkable that, in this study, most of the mutations detected in the evolved MA lines were also found in the RRDR region, including H442R (the most frequent one), D432G, S447L or S450L, mutations commonly found in *M. tuberculosis* rifampicin resistant isolates (*25*) and listed as drug resistance-conferring mutations by the WHO catalogue of *M. tuberculosis* mutations (*26*). The emergence of high-level resistance by RRDR *rpoB* mutations was the most frequent evolutionary pathway for rifampicin resistance during experimental evolution. The selection of high-level resistance mutations has occurred even under low rifampicin concentrations during this experimental evolution, as the appearance of drug resistance mutations could be selected by low antibiotic pressure (*66*). The emergence of a specific high-level resistance mutation, H442R, in a third of all Δ*nucS* lines, revealed an interesting phenomenon of evolutionary convergence, suggesting that multiple possible pathways to rifampicin resistance seem to be modulated and restricted to some preferred ones, as previously reported (*67*, *68*).

Additionally, different non-RRDR *rpoB* mutations and other *rpoB*-independent mutations conferring low or intermediate levels of resistance were also detected in evolved lines. Antibiotic resistance mutations associated with low and/or intermediate drug resistance are key intermediate steps in the pathways leading to high levels of resistance (*69*). No *rpoB* mutations in the MA lines were required to survive under subinhibitory concentrations of rifampicin. In fact, regulation of gene expression may support the growth of the MA lines under low antibiotic pressure, as rifampicin induces tolerance as an adaptative response to antibiotic stress in mycobacteria (*70*, *71*). Beyond this point, the acquisition of one or several rifampicin resistance-conferring mutations was required for survival and the wider mutational landscape of the Δ*nucS* strain was a strong advantage in the acquisition of resistance.

Since some drug resistant isolates lack *rpoB* mutation, alternative pathways for rifampicin resistance, involving different transporters and efflux pumps, antibiotic modifying enzymes and modulators of permeability, have been proposed (*12*). Several strategies have been used to screen for potential genes involved in drug resistance in mycobacteria, including the sequencing of resistant strains versus susceptible ones (*72*, *73*). Although antibiotic treatments often led to global changes in gene expression (*74*, *75*), very few genes enriched with mutations were found when bacterial cells were exposed to certain antibiotics (*16*, *18*). Here, we determined a subset of specific genes with high number of mutations under rifampicin selection, including some specific genes that could be associated with drug resistance and/or adaptative responses to the antibiotic. Some interesting candidates related with cell permeability (*trkB* and *mchK*) were validated as potential genes associated with low to intermediate levels of drug resistance, as previously reported (*30*, *31*). Further site-specific modification of the candidate genes would be required to confirm the association of specific mutations with rifampicin resistance.

In conclusion, MA evolution in combination with WGS constitutes an effective approach to study the total pool of genome-wide mutations upon antibiotic selection in mycobacteria. This strategy enabled us to evaluate the emergence of drug resistance mutations, the mechanisms and evolutionary pathways associated with drug resistance and the potential mutagenic effect of rifampicin. Finally, the appearance of genomic mutations under antibiotic selective pressure could be studied in pathogenic mycobacteria such as *M. tuberculosis*, to elucidate the impact of antibiotic treatments on the emergence and evolution of drug resistant isolates.

## Materials and Methods

### Bacterial strains and media

*M. smegmatis* wild type strain mc^2^ 155 (American Type Culture Collection, 700084) and its noncanonical MMR-deficient (Δ*nucS*) derivative (*21*) were grown at 37 °C in Middlebrook 7H9 broth or 7H10 agar (Difco) with 0.5% glycerol and Tween 80 (0.05% in 7H10 agar and 0.5% in 7H9 broth). The growth of bacterial strains in liquid medium was carried out in a shaker incubator at 250 rpm at 37°C.

### MA experiments

MA experiments were performed following a previously described procedure (*17*). However, in this study MA experiments were performed in the presence of increasing concentrations of rifampicin. In this work, 20 independent cell lines were generated from the wild type strain mc^2^ 155, and the same number from its isogenic Δ*nucS* derivative. These lines were independently evolved for 40 weeks (280 days) in parallel. Bacteria were initially streaked onto Middlebrook 7H10 plates supplemented with rifampicin at a starting concentration of 0.25 μg ml^−1^. Every week, a single random colony from each line was transferred to a new plate. The concentration of rifampicin was doubled every five weeks (five serial passages for each antibiotic concentration), ending the evolution at week 40 with a final concentration of 32 μg ml^−1^. The evolved lines generated at weeks 10, 20, 25, 30, 35 and 40 were stored at −80 °C for further analysis.

### Genomic DNA preparation and WGS analysis

Since most of the MA lines did not progress at the final rifampicin concentration of the evolution, we sequenced all evolved MA lines obtained at week 35 with 16 μg ml^−1^ rifampicin (38 lines: 18 wild type–derived lines and 20 Δ*nucS*-derived lines, numbered 1 to 20). The two parental strains (named mc^2^ and Δ*nucS* 0) were also sequenced as references. Genomic DNA was extracted from each of the 40 strains following the standard protocol for preparation of high-quality mycobacterial genomic DNA, as previously described (*76*). The integrity of each DNA sample and the absence of RNA contamination were confirmed by DNA agarose gel electrophoresis, while its concentration and purity were measured using a NanoDrop-2000 spectrophotometer (Thermo Fisher Scientific) and a Qubit 3.0 fluorometer (Life Technologies). WGS libraries were constructed with the NEBNext Ultra DNA Library Preparation Kit (Illumina, San Diego, CA). Sequencing was performed on the Illumina MiSeq instrument using a MiSeq v2 sequencing kit to obtain 250-bp paired-end reads.

### Bioinformatics analyses to detect SNPs and indels

WGS data were processed according to the following procedure: firstly, raw fastq files were filtered and trimmed with fastp (v0.12.5) using the following parameters (-- cut_by_quality3 --cut_window_size=10 --cut_mean_quality=20 --length_required=50 -- correction). Filtered fastq were mapped against the *M. smegmatis* mc^2^ 155 reference genome NC_018289.1 with the bwa mem algorithm (v0.7.12-r1039). Only reads that mapped the reference genome with a mapping quality equal to 60 were retained and duplicates were removed using picard tools (v. 2.18.0). Samtools (v.1.3.1) was used to obtain a pileup file, which was later scanned for genomic variants with VarScan2 (v2.3.7). For SNP calling we applied the following filters: minimum depth of 10 reads at the genomic position, presence of SNP in a minimum of 6 reads, minimum average quality 25, presence of SNP in both strands and at a minimum frequency of 0.5. Indels were called with gatk (v3.8-1-0-gf15c1c3ef), using the HaplotypeCaller program, and filtered with the VariantFiltration module the filters recommended for strict filtering: LowQd QD <2.0, HighFS FS >200.0, HighSOR SOR >3.0, Low40MQ MQ <40.0, LowReadPRS ReadPosRankSum 20.0, LowDepth DP <20. All detected variants were annotated with SnpEff (v.4.2). We analyzed the SNPs and indels detected in the evolved MA lines by comparison with their corresponding parental strain, wild type or Δ*nucS*, as appropriate. The set of variants found in the parental strains with respect to the *M. smegmatis* reference genome, detected in this study and in our previous work (*17*), were filtered using in-house Perl scripts and not considered for subsequent analyses. In this filtering step, SNPs having an indel in close proximity (10 nucleotides upstream or downstream) were removed. In the case of indels, those with low depth (DP<10) were also discarded from subsequent analyses.

### Calculation of mutation rates

Mutation rate per genome was calculated from WGS data by dividing the number of mutations by the total estimated generations (see Table 1), and mutation rate per nucleotide was calculated by dividing the number of mutations by the number of generations and the total sequenced sites in each genome (see Table 1 and Supplementary Tables S1 and S2). These calculations were performed for the overall mutations and for each specific type of change in the DNA. The number of generations was estimated as previously described (*17*). Firstly, we established a relationship between the number of viable cells present in a colony (*N*) and its area, as described by Lee *et al.*, 2012 (*77*). Then, wild type and Δ*nucS* MA lines were streaked on plates supplemented with each concentration used in the experimental evolution and the diameter of the colonies was measured after 7 days to estimate their number of viable cells. The total number of generations of each line was calculated as the sum of the generations per passage (*n*), being *n* = log_2_*N*. Statistical analyses (t-test) were performed to compare the mutation rates for each type of mutation in the presence and absence of rifampicin (Table S5).

### Sequencing of *rpoB* gene of the MA lines

To identify *rpoB* mutations, the full-length *rpoB* gene was sequenced from all the lines during experimental evolution (week 0, 10, 20, 25, 30, 35 and 40). In each case, *M. smegmatis rpoB* was amplified by PCR using the set of primers indicated in Supplementary Table S8. Five different overlapping PCR fragments were generated to cover the full-length *rpoB* gene in each sample. PCR products were sequenced using the Sanger method and the sequences were aligned with a *rpoB* wild type sequence to identify mutations.

### Sequencing of *trkB* and *mchK* genes of the MA lines

MA lines with mutations in *trkB* and *mchK* at week 35 of evolution were identified from WGS data. *trkB* and *mchK* genes were then sequenced in the same MA lines from the previous weeks to determine when each mutation emerged. The *trkB* and *mchK* genes were amplified by PCR using the primers indicated in Supplementary Table S8, and PCR products were sequenced using the Sanger method.

### Determination of rifampicin MICs

The rifampicin MICs of parental strains were evaluated by E-test on solid media. The rifampicin MICs of the MA lines stored at the different weeks of evolution were determined by broth microdilution method following the guidelines of the Clinical and Laboratory Standards Institute (CLSI) and using resazurin for the detection of mycobacterial growth as previously reported (*78*). Serial two-fold dilutions of rifampicin were performed in 100 μl of Middlebrook 7H9 broth in round-bottomed 96-well plates. All wells were then inoculated with 5 μl of overnight cultures of the *M. smegmatis* MA lines, previously diluted 1:100 (final dilution of approximately 1:2000). The plates were incubated at 37 °C for 48-72 h. Then, 30 μl of resazurin solution (200 μg ml^−1^) were added to all the wells, and plates were incubated overnight at 37 °C. The MIC was defined as the lowest drug concentration that prevented the change of the resazurin color from blue (oxidized form) to pink (reduced form), indicating the inhibition of the bacterial growth.

### Statistical analyses

Standard statistical analyses were performed using the *rstatix* package (v0.7.0) of R (v4.0.5).

## Data accessibility

Raw FASTQ files from WGS of this work, in presence of rifampicin, and the previous MA experiment in absence of antibiotic (*17*) are deposited into Sequence Read Archive (SRA) database with the BioProject accession numbers PRJNA899888 (this work) and PRJNA898800 (*17*).

## Acknowledgements

Editorial assistance was provided by Stuart L. Rulten.

## Supplementary Material

Supplementary Figure S1. Mutations in *trkB* and *mchK* in the MA lines.

Supplementary Table S1. Number of mutations and mutation rates of MA wild type lines, evolved under rifampicin selection.

Supplementary Table S2. Number of mutations and mutation rates of MA Δ*nucS* lines, evolved under rifampicin selection.

Supplementary Table S3. Number of mutations and mutation rates of MA wild type lines, evolved under antibiotic free conditions.

Supplementary Table S4. Number of mutations and mutation rates of MA Δ*nucS* lines, evolved under antibiotic free conditions.

Supplementary Table S5. Statistical analyses to compare mutation rates in the presence and absence of rifampicin of MA wild type and Δ*nucS* lines.

Supplementary Table S6. Mutations in *rpoB* gene and rifampicin MICs of the MA wild type lines of this study.

Supplementary Table S7. Mutations in *rpoB* gene and rifampicin MICs of the MA Δ*nucS* lines of this study.

Supplementary Table S8. Primers used in this study.

## References

1. O’Neill, J., Antimicrobial resistance: tackling a crisis for the health and wealth of nations. Rev. Antimicrob. Resist. (2014).

2. J. Davies, Inactivation of antibiotics and the dissemination of resistance genes. Science. 264, 375–382 (1994).

3. J. Blázquez, Hypermutation as a factor contributing to the acquisition of antimicrobial resistance. Clin. Infect. Dis. Off. Publ. Infect. Dis. Soc. Am. 37, 1201–1209 (2003).

4. S. M. Rosenberg, Evolving responsively: adaptive mutation. Nat. Rev. Genet. 2, 504–515 (2001).

5. M. A. Kohanski, M. A. DePristo, J. J. Collins, Sublethal antibiotic treatment leads to multidrug resistance via radical-induced mutagenesis. Mol. Cell. 37, 311–320 (2010).

6. J. Blázquez, J. Rodríguez-Beltrán, I. Matic, Antibiotic-induced genetic variation: how it arises and how it can be prevented. Annu. Rev. Microbiol. 72, 209–230 (2018).

7. B. Müller, S. Borrell, G. Rose, S. Gagneux, The heterogeneous evolution of multidrug-resistant *Mycobacterium tuberculosis*. Trends Genet. TIG. 29, 160–169 (2013).

8. World Health Organization, Global tuberculosis report 2021 (World Health Organization, Geneva, 2021; https://apps.who.int/iris/handle/10665/346387).

9. World Health Organization, Global tuberculosis report 2020 (World Health Organization, Geneva, 2020; https://apps.who.int/iris/handle/10665/336069).

10. E. A. Campbell, N. Korzheva, A. Mustaev, K. Murakami, S. Nair, A. Goldfarb, S. A. Darst, Structural mechanism for rifampicin inhibition of bacterial RNA polymerase. Cell. 104, 901–912 (2001).

11. M. Li, J. Lu, Y. Lu, T. Xiao, H. Liu, S. Lin, D. Xu, G. Li, X. Zhao, Z. Liu, L. Zhao, K. Wan, *rpoB* mutations and effects on rifampin resistance in *Mycobacterium tuberculosis*. Infect. Drug Resist. 14, 4119–4128 (2021).

12. G. Xu, H. Liu, X. Jia, X. Wang, P. Xu, Mechanisms and detection methods of *Mycobacterium tuberculosis* rifampicin resistance: the phenomenon of drug resistance is complex. Tuberc. Edinb. Scotl. 128, 102083 (2021).

13. G. Piccaro, D. Pietraforte, F. Giannoni, A. Mustazzolu, L. Fattorini, Rifampin induces hydroxyl radical formation in *Mycobacterium tuberculosis*. Antimicrob. Agents Chemother. 58, 7527–7533 (2014).

14. B. Van den Bergh, T. Swings, M. Fauvart, J. Michiels, Experimental design, population dynamics, and diversity in microbial experimental evolution. Microbiol. Mol. Biol. Rev. MMBR. 82, e00008–18 (2018).

15. J. E. Barrick, R. E. Lenski, Genome dynamics during experimental evolution. Nat. Rev. Genet. 14, 827–839 (2013).

16. H. Long, S. F. Miller, C. Strauss, C. Zhao, L. Cheng, Z. Ye, K. Griffin, R. Te, H. Lee, C.-C. Chen, M. Lynch, Antibiotic treatment enhances the genome-wide mutation rate of target cells. Proc. Natl. Acad. Sci. U. S. A. 113, E2498–2505 (2016).

17. A. Castañeda-García, I. Martín-Blecua, E. Cebrián-Sastre, A. Chiner-Oms, M. Torres-Puente, I. Comas, J. Blázquez, Specificity and mutagenesis bias of the mycobacterial alternative mismatch repair analyzed by mutation accumulation studies. Sci. Adv. 6, eaay4453 (2020).

18. H. O. Ozdemirel, D. Ulusal, S. Kucukyildirim Celik, Streptomycin and nalidixic acid elevate the spontaneous genome-wide mutation rate in *Escherichia coli*. Genetica. 149, 73–80 (2021).

19. A. Telenti, P. Imboden, F. Marchesi, D. Lowrie, S. Cole, M. J. Colston, L. Matter, K. Schopfer, T. Bodmer, Detection of rifampicin-resistance mutations in *Mycobacterium tuberculosis*. Lancet Lond. Engl. 341, 647–650 (1993).

20. B. P. Goldstein, Resistance to rifampicin: a review. J. Antibiot. (Tokyo). 67, 625–630 (2014).

21. A. Castañeda-García, A. I. Prieto, J. Rodríguez-Beltrán, N. Alonso, D. Cantillon, C. Costas, L. Pérez-Lago, E. D. Zegeye, M. Herranz, P. Plociński, T. Tonjum, D. García de Viedma, M. Paget, S. J. Waddell, A. M. Rojas, A. J. Doherty, J. Blázquez, A non-canonical mismatch repair pathway in prokaryotes. Nat. Commun. 8, 14246 (2017).

22. D. J. Jin, C. A. Gross, Mapping and sequencing of mutations in the *Escherichia coli rpoB* gene that lead to rifampicin resistance. J. Mol. Biol. 202, 45–58 (1988).

23. K. Severinov, M. Soushko, A. Goldfarb, V. Nikiforov, Rif^R^ mutations in the beginning of the *Escherichia coli rpoB* gene. Mol. Gen. Genet. MGG. 244, 120–126 (1994).

24. A. Koch, V. Mizrahi, D. F. Warner, The impact of drug resistance on *Mycobacterium tuberculosis* physiology: what can we learn from rifampicin? Emerg. Microbes Infect. 3, e17 (2014).

25. J. M. Musser, Antimicrobial agent resistance in mycobacteria: molecular genetic insights. Clin. Microbiol. Rev. 8, 496–514 (1995).

26. World Health Organization, Catalogue of mutations in Mycobacterium tuberculosis complex and their association with drug resistance (World Health Organization, Geneva, 2021; https://apps.who.int/iris/handle/10665/341981).

27. C. U. Köser, S. B. Georghiou, T. Schön, M. Salfinger, On the consequences of poorly defined breakpoints for rifampin susceptibility testing of *Mycobacterium tuberculosis* complex. J. Clin. Microbiol. 59, e02328–20 (2021).

28. World Health Organization, Technical report on critical concentrations for drug susceptibility testing of isoniazid and the rifamycins (rifampicin, rifabutin and rifapentine) (World Health Organization, Geneva, 2021; https://apps.who.int/iris/handle/10665/339275).

29. D. J. Wilson, The CRyPTIC Consortium, GenomegaMap: Within-species genome-wide dN/dS estimation from over 10,000 genomes. Mol. Biol. Evol. 37, 2450–2460 (2020).

30. A. Castañeda-García, T. T. Do, J. Blázquez, The K+ uptake regulator TrkA controls membrane potential, pH homeostasis and multidrug susceptibility in *Mycobacterium smegmatis*. J. Antimicrob. Chemother. 66, 1489–1498 (2011).

31. T. T. Do, J. Rodríguez-Beltran, E. Cebrián-Sastre, A. Rodríguez-Rojas, A. Castañeda-García, J. Blázquez, Inactivation of a new potassium channel increases rifampicin resistance and induces collateral sensitivity to hydrophilic antibiotics in *Mycobacterium smegmatis*. Antibiot. Basel Switz. 11, 509 (2022).

32. J. W. Schroeder, P. Yeesin, L. A. Simmons, J. D. Wang, Sources of spontaneous mutagenesis in bacteria. Crit. Rev. Biochem. Mol. Biol. 53, 29–48 (2018).

33. V. Katju, U. Bergthorsson, Old trade, new tricks: insights into the spontaneous mutation process from the partnering of classical mutation accumulation experiments with high-throughput genomic approaches. Genome Biol. Evol. 11, 136–165 (2019).

34. E. Denamur, O. Tenaillon, C. Deschamps, D. Skurnik, E. Ronco, J. L. Gaillard, B. Picard, C. Branger, I. Matic, Intermediate mutation frequencies favor evolution of multidrug resistance in *Escherichia coli*. Genetics. 171, 825–827 (2005).

35. D. I. Andersson, D. Hughes, Microbiological effects of sublethal levels of antibiotics. Nat. Rev. Microbiol. 12, 465–478 (2014).

36. L. Y. Song, M. Goff, C. Davidian, Z. Mao, M. London, K. Lam, M. Yung, J. H. Miller, Mutational consequences of ciprofloxacin in *Escherichia coli*. Antimicrob. Agents Chemother. 60, 6165–6172 (2016).

37. X. Giroux, W.-L. Su, M.-F. Bredeche, I. Matic, Maladaptive DNA repair is the ultimate contributor to the death of trimethoprim-treated cells under aerobic and anaerobic conditions. Proc. Natl. Acad. Sci. U. S. A. 114, 11512–11517 (2017).

38. M. N. Ragheb, M. K. Thomason, C. Hsu, P. Nugent, J. Gage, A. N. Samadpour, A. Kariisa, C. N. Merrikh, S. I. Miller, D. R. Sherman, H. Merrikh, Inhibiting the evolution of antibiotic resistance. Mol. Cell. 73, 157–165.e5 (2019).

39. J. W. Schroeder, T. S. Sankar, J. D. Wang, L. A. Simmons, The roles of replication-transcription conflict in mutagenesis and evolution of genome organization. PLoS Genet. 16, e1008987 (2020).

40. L. Laureti, I. Matic, A. Gutierrez, Bacterial responses and genome instability induced by subinhibitory concentrations of antibiotics. Antibiot. Basel Switz. 2, 100–114 (2013).

41. M. A. Kohanski, D. J. Dwyer, B. Hayete, C. A. Lawrence, J. J. Collins, A common mechanism of cellular death induced by bactericidal antibiotics. Cell. 130, 797–810 (2007).

42. J. J. Foti, B. Devadoss, J. A. Winkler, J. J. Collins, G. C. Walker, Oxidation of the guanine nucleotide pool underlies cell death by bactericidal antibiotics. Science. 336, 315–319 (2012).

43. D. J. Dwyer, P. A. Belenky, J. H. Yang, I. C. MacDonald, J. D. Martell, N. Takahashi, C. T. Y. Chan, M. A. Lobritz, D. Braff, E. G. Schwarz, J. D. Ye, M. Pati, M. Vercruysse, P. S. Ralifo, K. R. Allison, A. S. Khalil, A. Y. Ting, G. C. Walker, J. J. Collins, Antibiotics induce redox-related physiological alterations as part of their lethality. Proc. Natl. Acad. Sci. U. S. A. 111, E2100–2109 (2014).

44. J.-J. Kim, H.-M. Lee, D.-M. Shin, W. Kim, J.-M. Yuk, H. S. Jin, S.-H. Lee, G.-H. Cha, J.-M. Kim, Z.-W. Lee, S. J. Shin, H. Yoo, Y. K. Park, J. B. Park, J. Chung, T. Yoshimori, E.-K. Jo, Host cell autophagy activated by antibiotics is required for their effective antimycobacterial drug action. Cell Host Microbe. 11, 457–468 (2012).

45. R. R. Nair, D. Sharan, J. Sebastian, S. Swaminath, P. Ajitkumar, Heterogeneity of ROS levels in antibiotic-exposed mycobacterial subpopulations confers differential susceptibility. Microbiol. Read. Engl. 165, 668–682 (2019).

46. A. Paul, R. R. Nair, K. Jakkala, A. Pradhan, P. Ajitkumar, Elevated levels of three reactive oxygen species and Fe(II) in the antibiotic-surviving population of Mycobacteria facilitate *de novo* emergence of genetic resisters to antibiotics. Antimicrob. Agents Chemother. 66, e0228521 (2022).

47. Y. S. Patel, S. Mehra, Synergistic response of rifampicin with hydroperoxides on *Mycobacterium*: a mechanistic study. Front. Microbiol. 8, 2075 (2017).

48. F. Taddei, M. Radman, J. Maynard-Smith, B. Toupance, P. H. Gouyon, B. Godelle, Role of mutator alleles in adaptive evolution. Nature. 387, 700–702 (1997).

49. Y. Ram, L. Hadany, The evolution of stress-induced hypermutation in asexual populations. Evol. Int. J. Org. Evol. 66, 2315–2328 (2012).

50. R. C. MacLean, C. Torres-Barceló, R. Moxon, Evaluating evolutionary models of stress-induced mutagenesis in bacteria. Nat. Rev. Genet. 14, 221–227 (2013).

51. T. Swings, B. Van den Bergh, S. Wuyts, E. Oeyen, K. Voordeckers, K. J. Verstrepen, M. Fauvart, N. Verstraeten, J. Michiels, Adaptive tuning of mutation rates allows fast response to lethal stress in *Escherichia coli*. eLife. 6, e22939 (2017).

52. J. E. LeClerc, B. Li, W. L. Payne, T. A. Cebula, High mutation frequencies among *Escherichia coli* and *Salmonella* pathogens. Science. 274, 1208–1211 (1996).

53. A. Oliver, R. Cantón, P. Campo, F. Baquero, J. Blázquez, High frequency of hypermutable *Pseudomonas aeruginosa* in cystic fibrosis lung infection. Science. 288, 1251–1254 (2000).

54. A. Jolivet-Gougeon, B. Kovacs, S. Le Gall-David, H. Le Bars, L. Bousarghin, M. Bonnaure-Mallet, B. Lobel, F. Guillé, C.-J. Soussy, P. Tenke, Bacterial hypermutation: clinical implications. J. Med. Microbiol. 60, 563–573 (2011).

55. A. Giraud, I. Matic, M. Radman, M. Fons, F. Taddei, Mutator bacteria as a risk factor in treatment of infectious diseases. Antimicrob. Agents Chemother. 46, 863–865 (2002).

56. I. Chopra, A. J. O’Neill, K. Miller, The role of mutators in the emergence of antibiotic-resistant bacteria. Drug Resist. Updat. Rev. Comment. Antimicrob. Anticancer Chemother. 6, 137–145 (2003).

57. A. Giraud, I. Matic, O. Tenaillon, A. Clara, M. Radman, M. Fons, F. Taddei, Costs and benefits of high mutation rates: adaptive evolution of bacteria in the mouse gut. Science. 291, 2606–2608 (2001).

58. A. Giraud, M. Radman, I. Matic, F. Taddei, The rise and fall of mutator bacteria. Curr. Opin. Microbiol. 4, 582–585 (2001).

59. D. H. Mariam, Y. Mengistu, S. E. Hoffner, D. I. Andersson, Effect of *rpoB* mutations conferring rifampin resistance on fitness of *Mycobacterium tuberculosis*. Antimicrob. Agents Chemother. 48, 1289–1294 (2004).

60. S. Gagneux, C. D. Long, P. M. Small, T. Van, G. K. Schoolnik, B. J. M. Bohannan, The competitive cost of antibiotic resistance in *Mycobacterium tuberculosis*. Science. 312, 1944–1946 (2006).

61. I. Comas, S. Borrell, A. Roetzer, G. Rose, B. Malla, M. Kato-Maeda, J. Galagan, S. Niemann, S. Gagneux, Whole-genome sequencing of rifampicin-resistant *Mycobacterium tuberculosis* strains identifies compensatory mutations in RNA polymerase genes. Nat. Genet. 44, 106–110 (2011).

62. P. D. Sniegowski, P. J. Gerrish, R. E. Lenski, Evolution of high mutation rates in experimental populations of E. coli. Nature. 387, 703–705 (1997).

63. A. C. Shaver, P. G. Dombrowski, J. Y. Sweeney, T. Treis, R. M. Zappala, P. D. Sniegowski, Fitness evolution and the rise of mutator alleles in experimental *Escherichia coli* populations. Genetics. 162, 557–566 (2002).

64. J. E. Barrick, D. S. Yu, S. H. Yoon, H. Jeong, T. K. Oh, D. Schneider, R. E. Lenski, J. F. Kim, Genome evolution and adaptation in a long-term experiment with *Escherichia coli*. Nature. 461, 1243–1247 (2009).

65. V. Trinh, M.-F. Langelier, J. Archambault, B. Coulombe, Structural perspective on mutations affecting the function of multisubunit RNA polymerases. Microbiol. Mol. Biol. Rev. MMBR. 70, 12–36 (2006).

66. E. Gullberg, S. Cao, O. G. Berg, C. Ilbäck, L. Sandegren, D. Hughes, D. I. Andersson, Selection of resistant bacteria at very low antibiotic concentrations. PLoS Pathog. 7, e1002158 (2011).

67. D. Hughes, D. I. Andersson, Evolutionary trajectories to antibiotic resistance. Annu. Rev. Microbiol. 71, 579–596 (2017).

68. F. Baquero, J. L. Martínez, V. F Lanza, J. Rodríguez-Beltrán, J. C. Galán, A. San Millán, R. Cantón, T. M. Coque, Evolutionary pathways and trajectories in antibiotic resistance. Clin. Microbiol. Rev. 34, e0005019 (2021).

69. F. Baquero, Low-level antibacterial resistance: a gateway to clinical resistance. Drug Resist. Updat. Rev. Comment. Antimicrob. Anticancer Chemother. 4, 93–105 (2001).

70. J.-H. Zhu, B.-W. Wang, M. Pan, Y.-N. Zeng, H. Rego, B. Javid, Rifampicin can induce antibiotic tolerance in mycobacteria via paradoxical changes in *rpoB* transcription. Nat. Commun. 9, 4218 (2018).

71. S. N. Goossens, S. L. Sampson, A. Van Rie, Mechanisms of drug-induced tolerance in *Mycobacterium tuberculosis*. Clin. Microbiol. Rev. 34, e00141–20 (2020).

72. M. R. Farhat, B. J. Shapiro, K. J. Kieser, R. Sultana, K. R. Jacobson, T. C. Victor, R. M. Warren, E. M. Streicher, A. Calver, A. Sloutsky, D. Kaur, J. E. Posey, B. Plikaytis, M. R. Oggioni, J. L. Gardy, J. C. Johnston, M. Rodrigues, P. K. C. Tang, M. Kato-Maeda, M. L. Borowsky, B. Muddukrishna, B. N. Kreiswirth, N. Kurepina, J. Galagan, S. Gagneux, B. Birren, E. J. Rubin, E. S. Lander, P. C. Sabeti, M. Murray, Genomic analysis identifies targets of convergent positive selection in drug-resistant *Mycobacterium tuberculosis*. Nat. Genet. 45, 1183–1189 (2013).

73. H. Zhang, D. Li, L. Zhao, J. Fleming, N. Lin, T. Wang, Z. Liu, C. Li, N. Galwey, J. Deng, Y. Zhou, Y. Zhu, Y. Gao, T. Wang, S. Wang, Y. Huang, M. Wang, Q. Zhong, L. Zhou, T. Chen, J. Zhou, R. Yang, G. Zhu, H. Hang, J. Zhang, F. Li, K. Wan, J. Wang, X.-E. Zhang, L. Bi, Genome sequencing of 161 *Mycobacterium tuberculosis* isolates from China identifies genes and intergenic regions associated with drug resistance. Nat. Genet. 45, 1255–1260 (2013).

74. H. I. M. Boshoff, T. G. Myers, B. R. Copp, M. R. McNeil, M. A. Wilson, C. E. Barry, The transcriptional responses of *Mycobacterium tuberculosis* to inhibitors of metabolism: novel insights into drug mechanisms of action. J. Biol. Chem. 279, 40174–40184 (2004).

75. J. Briffotaux, S. Liu, B. Gicquel, Genome-wide transcriptional responses of *Mycobacterium* to antibiotics. Front. Microbiol. 10, 249 (2019).

76. J. T. Belisle, S. B. Mahaffey, P. J. Hill, “Isolation of *Mycobacterium* species genomic DNA” in *Mycobacteria Protocols: Second Edition*, T. Parish, A. C. Brown, Eds. (Humana Press, Totowa, NJ, 2009), Methods in Molecular Biology, pp. 1–12.

77. H. Lee, E. Popodi, H. Tang, P. L. Foster, Rate and molecular spectrum of spontaneous mutations in the bacterium *Escherichia coli* as determined by whole-genome sequencing. Proc. Natl. Acad. Sci. U. S. A. 109, E2774–2783 (2012).

78. J.-C. Palomino, A. Martin, M. Camacho, H. Guerra, J. Swings, F. Portaels, Resazurin microtiter assay plate: simple and inexpensive method for detection of drug resistance in *Mycobacterium tuberculosis*. Antimicrob. Agents Chemother. 46, 2720–2722 (2002).

