## Supplementary Figure S1 for "Selective pressure by rifampicin modulates mutation rates and evolutionary trajectories of mycobacterial genomes"

**A**

| *trkB* mutations | | | | | | |
|---|---|---|---|---|---|---|
| Week | 10 | 20 | 25 | 30 | 35 | 40 |
| [Rif] (µg/ml) | 0.5 | 2 | 4 | 8 | 16 | 32 |
| Line | | | | | | |
| Δ*nucS* 5 | A130A | A130A | A130A | A130A | A130A | A130A |
| Δ*nucS* 12 | NO | NO | NO | NO | W111R | extinct |
| Δ*nucS* 18 | NO | V91A | V91A | V91A | V91A | V91A |
| Δ*nucS* 20 | NO | NO | NO | G12D | G12D | G12D |

**B**

| *mchK* mutations | | | | | | |
|---|---|---|---|---|---|---|
| Week | 10 | 20 | 25 | 30 | 35 | 40 |
| [Rif] (µg/ml) | 0.5 | 2 | 4 | 8 | 16 | 32 |
| Line | | | | | | |
| $mc^2$ 4 | NO | NO | NO | NO | Δ - 1c | extinct |
| $mc^2$ 6 | NO | NO | NO | T163I | T163I | extinct |
| $mc^2$ 19 | NO | NO | NO | A216V | A216V | extinct |
| Δ*nucS* 1 | NO | NO | NO | R120Q | R120Q | R120Q |
| Δ*nucS* 2 | NO | NO | NO | NO | E316G | E316G |
| Δ*nucS* 8 | NO | NO | NO | P321L | P321L | extinct |
| Δ*nucS* 9 | NO | NO | V203A | V203A | V203A | V203A |
| Δ*nucS* 19 | NO | NO | Q265R | Q265R | Q265R | Q265R |

Rif MIC (µg $ml^{-1}$) — 1 - 2 No resistant; 4 - 16 Low; 32 - 128 Intermediate; 256 - 1024 High

**Figure S1. Mutations in *trkB* and *mchK* in the MA lines. A**. Emergence of *trkB* mutations in the MA lines. **B**. Emergence of *mchK* mutations in the MA lines. Tables show the appearance of mutations in *trkB* and *mchK* in each MA line during experimental evolution (in weeks). Only MA lines with mutations in these genes are represented. Levels of rifampicin resistance of the evolved lines are indicated according to their MICs values, with the following color code: no resistance, 1 – 2 µg $ml^{-1}$ (grey); low, 4 – 16 µg $ml^{-1}$ (light red); intermediate, 32 – 128 µg $ml^{-1}$ (medium red); and high, 256 – 1024 µg $ml^{-1}$ (dark red).
