## Supplementary Table S8 for "Selective pressure by rifampicin modulates mutation rates and evolutionary trajectories of mycobacterial genomes"

**Table S8. Primers used in this study**

| Primer name* | Sequence | Target gene | Source |
| --- | --- | --- | --- |
| M.smegmatis-rpoB-FWD (f) | GCTGATCCAGAACCAGATCC | *rpoB* (MSMEI_1328, MSMEG_1367) | (*16*) |
| M.smegmatis-rpoB-REV (r) | GATGACACCGGTCTTGTCG | *rpoB* (MSMEI_1328, MSMEG_1367) | (*16*) |
| rpoB-F1 (f) | TGTCGTTGCGTCCAGGGTTCTGGA | *rpoB* (MSMEI_1328, MSMEG_1367) | This study |
| rpoB-R1 (r) | GCTCCACGATCTGCTCGTTGGTCC | *rpoB* (MSMEI_1328, MSMEG_1367) | This study |
| rpoB-F2 (f) | GTATCGACCGCAAGCGCCGCCAGC | *rpoB* (MSMEI_1328, MSMEG_1367) | This study |
| rpoB-R2 (r) | CGAACCGATCAGACCGATGTTGGG | *rpoB* (MSMEI_1328, MSMEG_1367) | This study |
| rpoB-F3 (f) | GACGTGCACCCCAGCCACTACGGC | *rpoB* (MSMEI_1328, MSMEG_1367) | This study |
| rpoB-R3 (r) | AGCTTGGTGTCGCGGGCATCGATC | *rpoB* (MSMEI_1328, MSMEG_1367) | This study |
| rpoB-F4 (f) | GAACCGCCTGGTCGAAGAGGACGT | *rpoB* (MSMEI_1328, MSMEG_1367) | This study |
| rpoB-R4 (r) | ACATGTAGCCAACCGTCACCGGGT | *rpoB* (MSMEI_1328, MSMEG_1367) | This study |
| rpoB-F5 (f) | ACGCCGACGGCAAGGCGACGCTGT | *rpoB* (MSMEI_1328, MSMEG_1367) | This study |
| rpoB-R5 (r) | CTACGCGAGATCCTCGACGGACGC | *rpoB* (MSMEI_1328, MSMEG_1367) | This study |
| TrkB2769seqF (f) | ATGAAAGTCGCCATCGCCGGTGCC | *trkB* (MSMEI_2701, MSMEG_2769) | This study |
| TrkB2769seqR (r) | CTAACGCCGCGTGGGCCGCAGCAG | *trkB* (MSMEI_2701, MSMEG_2769) | This study |
| MchK1945seqF (f) | GTGGCTAAAGGCAGGTTACGGCGC | *mchK* (MSMEI_1903, MSMEG_1945) | This study |
| MchK1945seqR (r) | TCATCGTTCGGCGTCCGCACTGCG | *mchK* (MSMEI_1903, MSMEG_1945) | This study |

* (f): forward, (r): reverse.
